# Adult lifespan effects on functional specialization along the hippocampal long axis

**DOI:** 10.1101/2024.10.04.616732

**Authors:** Caitlin R. Bowman, Cara I. Charles, Saisha M. Birr

## Abstract

There has been increasing attention to differences in function along the hippocampal long axis, with the posterior hippocampus proposed to be well suited to representing fine-grained details and the anterior hippocampus having coarser representations. Whether long axis functional specialization persists into older age is not well understood, despite known age-related declines in the level of detail in memories. In this study, we used a large database of resting state fMRI data (n =322 humans of both sexes included) from across the adult lifespan (ages 18-88) to determine the degree of functional differentiation across the hippocampal posterior-anterior axis. Our first approach was to measure the similarity among signals within hippocampal subregions. We found that signals within the most posterior hippocampal subregion became more similar to one another in older age, but this effect did not relate to memory performance. As a second approach, we measured functional connectivity between hippocampal subregions and the rest of the brain. The functional connectivity profiles of the posterior and anterior hippocampal subregions became more distinct from one another with increasing age, driven by relative stability in anterior hippocampal functional connectivity but more significant age-related differences in posterior subregions of the hippocampus. Stronger connectivity between the anterior hippocampus and several cortical regions was associated with better memory performance in older adults, suggesting that maintenance of anterior hippocampal connectivity may help preserve episodic memory among high performing older adults.

## 1. Introduction

The hippocampus is critical for episodic memory function (Scoville & Milner, 1957) and shows structural and functional decline in healthy (Hu & Li, 2020; Leal & Yassa, 2015; O’Shea et al., 2016) and pathological aging (Anand & Dhikav, 2012; Peng et al., 2014). A functional differentiation along the hippocampal long axis has been proposed based on work in healthy young adults and in animals, with posterior hippocampus representing fine-grained information and anterior hippocampus having coarser representations that integrate across larger temporal and spatial windows (Brunec et al., 2018; Poppenk et al., 2013). This long axis differentiation may emerge due to differences in the granularity of signals within the hippocampus. A prior study in young adults showed less correlated signals across voxels within the posterior hippocampus compared to those within the anterior hippocampus (Brunec et al., 2018). The authors took this as evidence that the posterior hippocampus is well suited to encoding information over narrow spatial and temporal windows, because its more variable and dynamic signals would be capable of capturing idiosyncratic, fine-grained details. However, a later study found a different patten: signals were least similar to one another in intermediate portions of the hippocampus and more similar in the furthest anterior and posterior portions (Thorp et al., 2022). Thus, there remains some question of the nature of intra-hippocampal signals across the hippocampal long axis. A prior aging study showed increases in the similarity of signals within the posterior medial temporal lobe, including the posterior hippocampus, and that these increases were associated with poorer episodic memory abilities (Salami et al., 2016), providing initial evidence that disruptions to intra-hippocampal signals contribute to age-related declines in episodic memory.

Patterns of structural and functional connectivity between the hippocampus and the rest of the brain may also contribute to hippocampal long axis specialization (Catenoix et al., 2011; Duvernoy et al., 2005; Fanselow & Dong, 2010; L. E. Frank et al., 2019; Jung et al., 1994; Kahn et al., 2008). Prior work in healthy young adults has shown that parts of lateral prefrontal cortex and lateral parietal cortex have stronger functional connectivity with the posterior than the anterior hippocampus, while the ventromedial prefrontal cortex showed greater coupling with the anterior compared to posterior hippocampus (L. E. Frank et al., 2019). Other work has positioned the hippocampal functional gradient within larger medial temporal lobe (MTL) systems (Libby et al., 2012; Ranganath & Ritchey, 2012; Rugg & Vilberg, 2013). According to this framework, the posterior hippocampus is part of the posterior medial system that includes the parahippocampal cortex, retrosplenial cortex, posterior cingulate, precuneus, angular gyrus, and anterior thalamus. The anterior hippocampus is part of the anterior temporal system, which includes the perirhinal cortex, lateral orbitofrontal cortex, amygdala, and temporal pole. In older adults, separable posterior medial and anterior temporal networks have been detected (Das et al., 2015), and healthy aging has been associated with weaker within-network connections for both the posterior medial and anterior temporal networks (Hrybouski et al., 2023). A prior study that included older adults in a sample that was predominantly children, young, and middle aged adults showed age-related decreases in connectivity between the posterior hippocampus and the anterior cingulate and medial superior frontal gyrus (Panitz et al., 2021). Yet, the extent to which these effects are driven by differences in advanced age and their relationship to memory abilities remains unclear.

In the present study, we tested the degree to which patterns of hippocampal functional specialization persist across the adult lifespan and how any age-related differences in hippocampal specialization contribute to known age-related declines in episodic memory (Fraundorf et al., 2019; Old & Naveh-Benjamin, 2008; Spencer & Raz, 1995; Toner et al., 2009). We used resting-state fMRI data from individuals aged 18-88 (n = 322) from the Cambridge Centre for Ageing and Neuroscience (CamCAN) dataset (Shafto et al., 2014). We compared signals across three anatomically defined segments along the hippocampal posterior-anterior axis (tail, body, head) in terms of both the similarity of signals within each hippocampal subregion and functional connectivity between each subregion and the rest of the brain. We related these metrics to age-related differences in performance on an episodic memory task. Based on the proposed role of the posterior hippocampus in representing fine-grained details in memory and known age-related decline in episodic memory, we expected to find disproportionate age-related differences in posterior hippocampal function.

## 2. Methods

### 2.1. Data and Code Availability

The raw data come from a publicly available dataset (https://www.cam-can.org). Intervoxel similarity values (IVS), functional connectivity values, scores on the behavioral tasks used, and nuisance covariates for individual subjects are publicly available through the Open Science Framework (https://osf.io/qba8n/). Analytic code is also available.

### 2.2. Participants

All data came from the publicly available CamCAN dataset (Shafto et al., 2014). From the larger dataset, we obtained resting state fMRI scans, anatomical MRI scans, and behavioral data from 653 individuals aged 18-88. To ascertain cognitive health, all participants underwent cognitive testing with the Mini-Mental State Exam (MMSE). Those who scored >24/30 points and who did not report any neurological conditions were invited to participate in the fMRI portion of the study. Of the 653 participants obtained from the database, we excluded one subject due to missing functional data and two others due to failed parcellation of their anatomical images. We excluded 24 subjects because they had fewer than 10 voxels in one or more hippocampal subregion. We also excluded for excessive motion. Our threshold for complete exclusion was having any single framewise displacement value greater than or equal to 1 mm or having fewer than 5 minutes of timepoints included after scrubbing (details below). These strict motion exclusions are necessary because functional connectivity analyses of rest data are especially sensitive to motion (Power et al., 2012, 2014). This procedure led to an additional 270 participants excluded. Thirty-five additional participants were excluded for having both excessive motion and too few voxels in a hippocampal subregion. The final sample for analyses involving only the neuroimaging data was 322 subjects. Because the emotional memory task that served as the best measure of episodic memory abilities was only administered to half of the participants in the original CamCAN study, a subset of the sample used for neuroimaging analyses was carried forward to analyses linking brain indices to behavior. The sample for brain-behavior analyses included 156 subjects. For further details about the distribution of age, sex, education and MMSE scores in these samples, see Table 1.

**Table 1:** Sample characteristics separated by decades-based age groups.

| fMRI data only analyses |  |  |  |  | Brain-behavior analyses |  |  |  |
| --- | --- | --- | --- | --- | --- | --- | --- | --- |
| Age Group | N included | % Female | Mean MMSE | Mean Years Education | N included | % Female | Mean MMSE | Mean Years Education |
| 18-29 | 51 | 55% | 29.2 | 15.3 | 26 | 46% | 29.4 | 15.4 |
| 30-39 | 66 | 53% | 28.7 | 16.0 | 31 | 48% | 29.3 | 15.8 |
| 40-49 | 71 | 46% | 29.2 | 15.8 | 33 | 55% | 29.3 | 15.9 |
| 50-59 | 43 | 63% | 29.4 | 15.5 | 24 | 62% | 29.6 | 16.0 |
| 60-69 | 45 | 42% | 28.9 | 14.2 | 19 | 42% | 28.8 | 13.9 |
| 70+ | 46 | 33% | 28.1 | 14.2 | 23 | 30% | 28.1 | 13.7 |
| All | 322 | 47% | 29.0 | 15.8 | 156 | 48% | 29.1 | 15.3 |

### 2.3. Data acquisition and processing

Participants in the Cam-CAN study completed a series of behavioral tasks across a Stage 1 interview phase. They then completed a Stage 2 phase involving detailed cognitive testing and MRI. Testing occurred over three sessions (Shafto et al., 2014).

#### 2.3.1. MRI Data Acquisition

Full details of the MRI acquisition protocol have been previously reported (Shafto et al., 2014). From the larger dataset, we used the resting state functional run and the T1-weighted and T2-weighted structural images. During the resting state functional run, participants were asked to keep their eyes closed while 261 EPI volumes were acquired (1.97 second TR, 3 mm x 3 mm x 4.44 mm voxel), with a total acquisition time of 8 minutes and 40 seconds.

#### 2.3.2. Anatomical data preprocessing and regions of interest selection

We defined anatomical regions of interest in each subject’s native space from Freesurfer 7.1.1 (Fischl, 2012). We applied Freesurfer’s standard cortical parcellation and subcortical segmentation to the T1-weighted anatomical image. Afterwards, we further segmented the hippocampus by applying Freesurfer’s segmentHA_T2 protocol using both the T1-weighted and T2-weighted anatomical images as inputs (Iglesias et al., 2015). Freesurfer’s automated hippocampal segmentation was developed from manual segmentations of ultra-high resolution ex vivo data and high-resolution in vivo data with a protocol developed from several sources (De Nó, 1934; Duvernoy et al., 2013; Green & Mesulam, 1988; Insausti & Amaral, 2012; Rosene & Van Hoesen, 1987; Van Leemput et al., 2009). Prior work has shown good to excellent reliability of these segmentations (Kahhale et al., 2023). From the hippocampal segmentation, we selected the tail, body, and head (separately for the right and left hemispheres) as our posterior, intermediate, and anterior hippocampal regions of interest (ROIs), respectively (Figure 1). In this segmentation protocol, the boundary between the tail and the body is defined by the point where the fornix fully connects to the hippocampus (Iglesias et al., 2015). The boundary between the body and the head is defined by the caudal boundary of the uncus (Van Leemput et al., 2009). Because some findings have suggested that Freesurfer’s overall hippocampal segmentation may perform differently across age groups (Srinivasan et al., 2020), we verified that our findings held when using a multi-atlas segmentation protocol to define the hippocampus (Doshi et al., 2016) and co-registered the tail, body, and head ROIs from Freesurfer into that hippocampal space. The functional hippocampal signals were similar regardless of the segmentation protocol used, so we retained the Freesurfer segmentation for simplicity.

**Figure 1.**
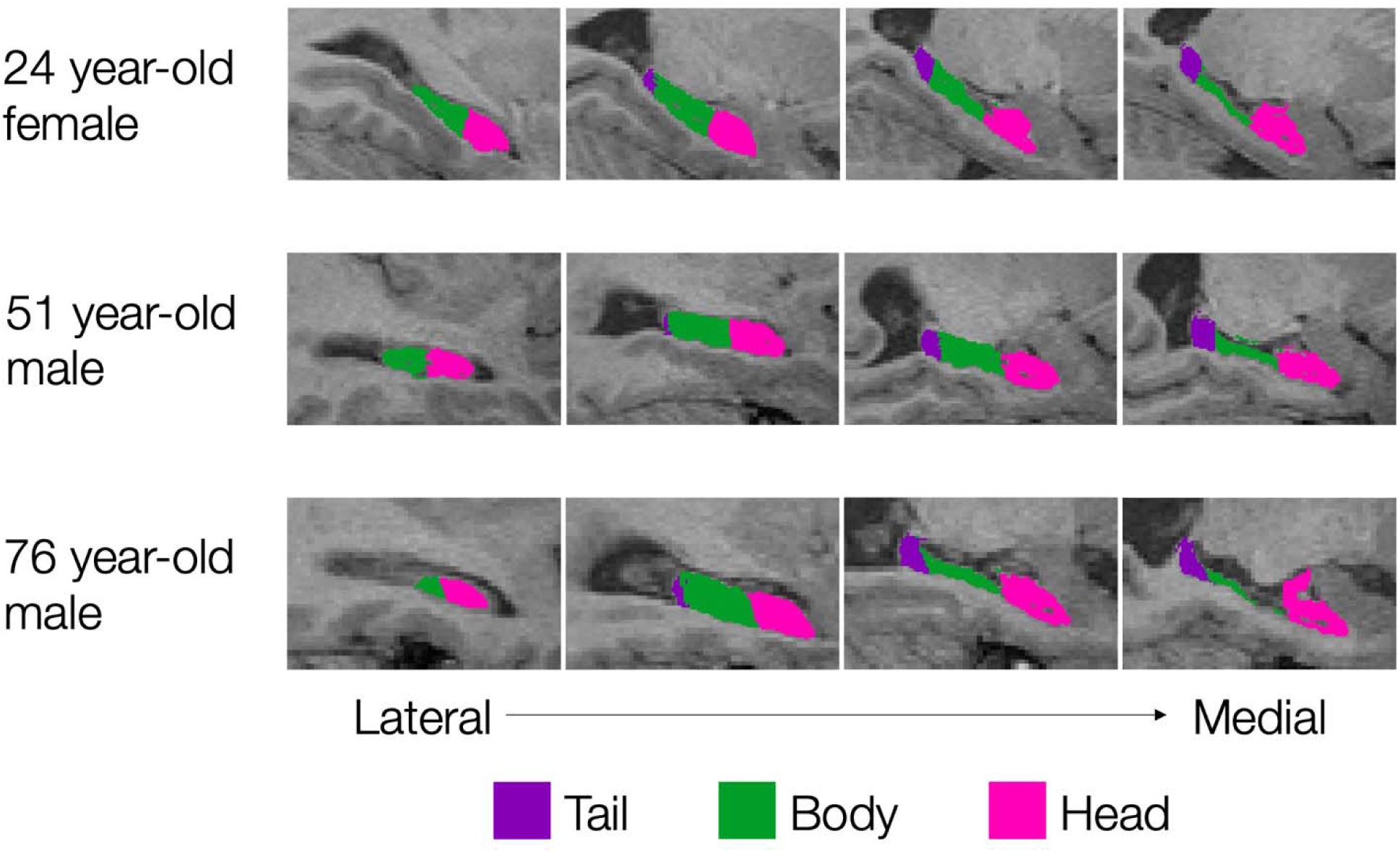
Hippocampal subregions of interest. The hippocampal subregions from one randomly selected young adult, one middle aged adult, and one older adult are depicted on each subject’s structural scan. The hippocampus was divided into three segments with the tail (purple) serving as the most posterior region, the body (green) serving as an intermediate region, and the head (pink) serving as the most anterior region. Boundaries between regions were determined by Freesurfer’s hippocampal segmentation protocol and all analyses were computed in native space.

We also used the cortical parcellation and subcortical segmentation from Freesurfer to define anatomical regions of interest to serve as targets in the functional connectivity analysis. We used 42 distinct anatomical labels (84 total regions across left and right hemispheres) from the Desikan-Killiany cortical atlas and the subcortical segmentation (excluding the hippocampus since it served as the seed). For a full list of target regions, see Supplementary Table S1. Compared to a whole brain, voxelwise approach, this approach averages across many voxels to increase the reliability of the signal in each ROI and reduces the number of pairwise connections to correct for when multiple comparison corrections are needed. Using a whole-brain ROI approach also aids in interpreting findings by allowing us to leverage the known structural and functional properties of these regions (Ritchey & Cooper, 2020). After defining ROIs from the structural images, we used the mutual information algorithm from Advanced Normalization Tools (ANTS) version 2.1.0 to register and transform the anatomical images to functional space.

#### 2.3.3. fMRI preprocessing

We first stripped the skull from each subject’s resting state functional run using the Brain Extraction Tool from FSL (Jenkinson et al., 2006), then used FSL’s MCFLIRT to motion correct the functional data, realigning all volumes to the middle volume. Using FEAT (fMRI Expert Analysis Tool) in FSL, the brain extracted, and motion corrected functional images were subjected to a high-pass temporal filter (100 sec) to remove low frequency drift in the scanner signal. Functional images were then minimally spatially smoothed using a 2-mm FWHM kernel.

Because functional connectivity measures can be especially sensitive to motion (Murphy et al., 2013; Power et al., 2012), we used the fMRI quality assurance tool (https://github.com/poldrack/fmriqa) to identify subjects who moved excessively as well as individual timepoints with excessive motion. We computed framewise displacement (FD) scores, the temporal derivative of the signal variance over voxels (DVARS), translational and rotational motion, and the temporal derivatives of translational and rotational motion. We excluded all data from any subject who had any single FD value > 1 mm. For the remaining subjects, we identified individual timepoints with movement FD > .5 mm or DVARS > .5% and removed those timepoints as well as the timepoint immediately prior and two timepoints immediately after each motion-flagged timepoint. Subjects for whom fewer than 42% of their timepoints were usable (i.e., 5 minutes of data included) were excluded entirely. We used the 12 translational and rotational motion parameters, plus the mean signals from white matter, CSF, and whole brain and their temporal derivatives as volume-by-volume motion covariates when computing inter-voxel similarity within hippocampal subregions and functional connectivity between hippocampal subregions and the rest of the brain.

#### 2.3.4. Inter-voxel similarity

Following pre-processing, we used the fslmeants function from FSL to extract a time course from each voxel within each hippocampal subregion separately from the right and left hemisphere of each subject. We then used custom R code (R Core Team, 2021) to compute the partial correlation between the timeseries of each pair of voxels in each subregion, controlling for volume-by-volume motion estimates. Following a Fisher’s Z transformation of the resulting r-values, we averaged across the pairwise z-values for each subregion, leading to six inter-voxel similarity values for each subject (3 hippocampal subregions x 2 hemispheres).

#### 2.3.5. Functional connectivity

We used the fslmeants function from FSL to extract a mean time course from each of the 84 target regions of interest (42 labels x right and left hemisphere). We then calculated the functional connectivity scores as a partial correlation between each hippocampal seed region (right and left hippocampal head, body, and tail = 6 seed regions) and each target region (right and left of 42 cortical and subcortical targets = 84 target regions), controlling for volume-by-volume motion estimates. This procedure led to 504 pairwise connections for each subject. We then Fisher’s Z transformed the resulting r-values and submitted these connectivity values to group-level analyses.

#### 2.3.6. Episodic memory performance

From the cognitive measures obtained, only the emotional memory task specifically tested episodic memory, and we therefore used it as our episodic memory measure. However, only half of the participants in the study were administered this task (the other half did an emotion regulation task instead). This led to 156 individuals who were included in imaging analyses and had episodic memory scores available.

In the emotional memory task, participants first underwent a study phase where they were shown a background picture for 2 seconds before an object would appear superimposed on the background. Participants were instructed to create a story linking the object to the background. After 8 seconds, the screen would advance to the next trial. There were 120 of these trials in the study phase, and participants were not told that there would be a memory test. The emotional component of the task came from the background pictures, which were from the International Affective Pictures Set (IAPS) (Lang et al., 1999). The background picture could depict a positive situation, a neutral situation, or a negative situation. There was a 10-minute retention interval between the study and test phases. There were three components of each test trial, testing different aspects of memory. First, as a measure of perceptual priming, participants saw a degraded version of an object and were asked to identify it. Next, as a measure of object recognition, they saw the full, non-degraded object image and were asked whether it was old (presented in the study phase) or new, also indicating their confidence. Lastly, as a measure of associative and contextual memory, participants were asked the valence (positive, neutral, or negative) of the background image presented with the object. There were 120 trials with old, studied objects and 40 trials with new objects. In our analyses, we included the object recognition and background valence memory as indices of episodic memory. We averaged across the valence conditions. For object recognition, we computed the average hit rate (collapsed across confidence levels) across the three valence conditions, and we subtracted the single false alarm rate (Figure 2A). For the background valence memory, we computed the averaged hit rate across the three valence conditions and the average false alarm rate (e.g., a positive valence false alarm would be responding ‘positive’ to an object associated with a neutral or negative background image). We then computed the difference between the hit and false alarm rates (Figure 2B). These two metrics were highly correlated with one another *r*(154) = .72, *p* < .001, so we created an episodic memory composite score by calculating z-scores for each measure, then averaged across the two resulting z-scores for each participant (Figure 2C).

**Figure 2.**
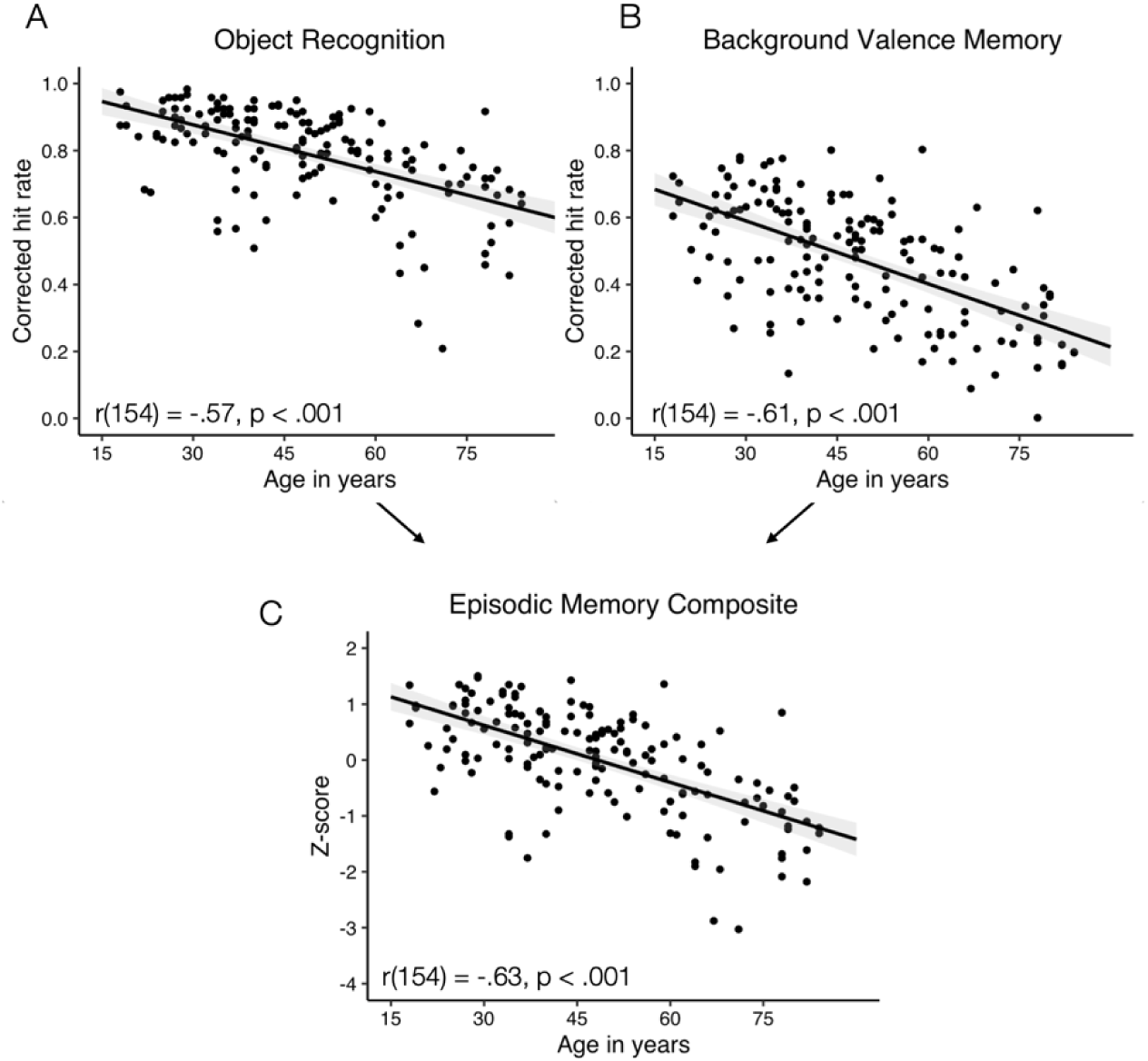
Relationship between age, episodic memory performance, and visual short-term memory. The relationship between age (in years) and the corrected hit rate (hit rate – false alarm rate) for object recognition (A) and background picture valence (B) from the episodic memory task. (C) The relationship between age and the episodic memory composite derived from the object-emotional background task. The y-axis for the composite is the mean z-score across the two measures included in the composite (object recognition, and background valence memory). In A-C, r values represent the zero-order correlation between age and the given behavioral metric.

### 2.4. Statistical analyses

Group-level statistical analyses were computed in R (R Core Team, 2021) and the full analytic code is available in an OSF repository (https://osf.io/qba8n/). In all analyses that compared across ages, we included participant gender (dummy coded as female = 0, male = 1), subject-level motion (defined as the proportion of excluded timepoints), and overall bilateral hippocampal volume (measured as the proportion of total intracranial volume) as covariates unless otherwise noted. The motion and hippocampal volume covariates were not significantly related to one another across the entire sample, *r*(320) = .029, *p* = .609, or in either the female, *r*(151) = -.011, *p* = .892, or male, *r*(167) = .052, *p* = .499, portions of the sample. Thus, we did not see signs of collinearity among our covariates, and we controlled for these other factors when determining age effects in hippocampal function. Linear mixed effects models were computed using the nlme package in R (Pinheiro et al., 2025). To account for potential heteroskedasticity in the data, we computed robust standard errors for all regression models using the Sandwich package version 3.1.1 in R (Zeileis et al., 2020). When computing multiple tests on the same data, we used the Holm-Bonferroni method to correct for multiple comparisons.

## 3. Results

### 3.1. Age-related differences in within hippocampal signals

Our first approach to understanding the impact of age on hippocampal functional specialization was to compare signals from within each of the three hippocampal regions. As in prior studies, we computed the correlation across timeseries for each pair of voxels within each hippocampal region (‘inter-voxel similarity’) (Brunec et al., 2018; Thorp et al., 2022). Higher values on this metric indicate that the voxels within a given hippocampal subregion have more similar signals to one another, with higher similarity proposed to subserve integration across memories and generalization as opposed to specificity and discrimination (Bowman & Zeithamova, 2018; Brunec et al., 2018; Collin et al., 2015; Poppenk et al., 2013; Schlichting et al., 2015). The group-level inter-voxel similarity values across all ages are depicted in Figure 3A. A linear mixed effects model indicated a significant effect of hippocampal region on inter-voxel similarity, ß = -.016, *SE* = .005, *t* = -3.19, *p* = .001, with the hippocampal tail having significantly *higher* inter-voxel similarity than both the hippocampal body, *t*(321) = 6.89, *p* < .001, *d* = 1.54, and hippocampal head, *t*(321) = 12.22, *p* < .001, *d* = 2.88, and the hippocampal body having higher inter-voxel similarity than the hippocampal head, *t*(321) = 7.15, *p* < .001, *d* = 1.55. There was also a main effect of hemisphere, *ß* = -.032, *SE* = .015, *t* = -2.073, *p* = .038, with higher inter-voxel similarity in the right hemisphere (M = .14, SD = .03) compared to the left hemisphere (M = .13, SD = .03). There were also significant age x hemisphere, *ß* = .0007, SE = .0003, *t* = 2.428, *p* = .015, region x hemisphere, *ß* = .017, *SE* = .007, *t* = 2.41, *p* = .016, and age x region x hemisphere interaction effects, *ß* = -.0003, *SE* = .0001, *t* = -2.184, *p* = .029. Figure 3B presents the inter-voxel similarity values for the left and right hippocampus separated into six decades-based age groups. Visual inspection suggests that the general tail > body > head pattern was apparent across age groups with the exception of the 18–29-year-old group in the right hippocampus, which showed the highest inter-voxel similarity in the body of the hippocampus. In addition, the pattern of higher inter-voxel similarity in the hippocampal tail vs body and head became more pronounced in older age groups.

**Figure 3.**
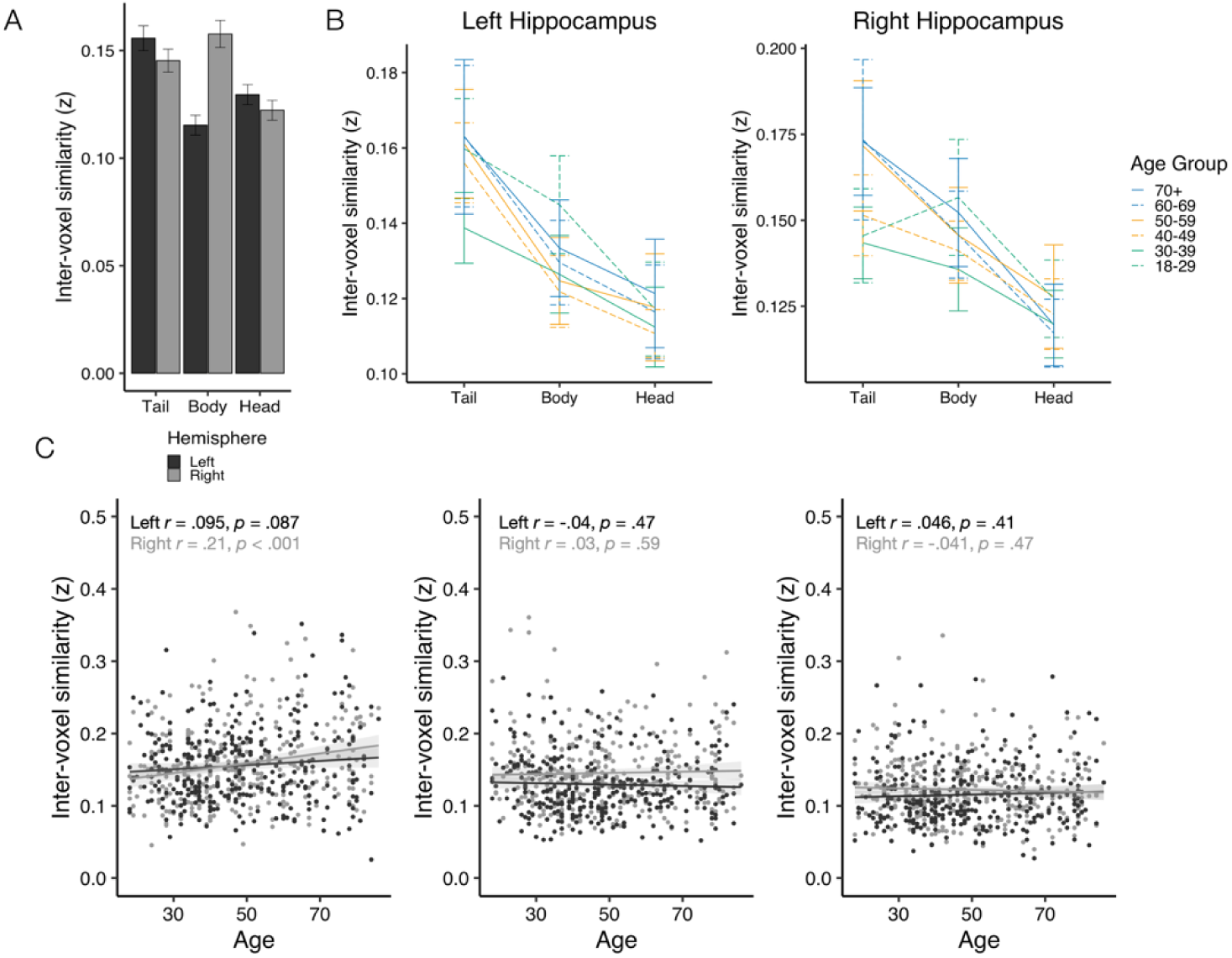
Age-related differences in the correlation among intra-hippocampal signals. (A) Mean inter-voxel similarity across all ages separated by hippocampal subregion and left vs. right hemisphere. (B) Mean intervoxel similarity values for the left and right hippocampal subregions separated by approximately decades-based age groups. The relationship between continuous age (in years) and inter-voxel similarity in each hippocampal subregion. Trendlines represent the zero-order relationship between age and intervoxel similarity. In A-C, error bars/shaded region represent 95% confidence intervals.

To quantify patterns of age-related differences in the intra-hippocampal subregion signal similarity, we examined the relationship between age and inter-voxel similarity for each hippocampal subregion separately for the left and right hemisphere (Figure 3C). We conducted multiple linear regressions for each hippocampal subregion using age to predict intervoxel similarity, controlling for gender, motion, and hippocampal volume. Results showed a significant positive relationship between age and IVS in the right hippocampal tail, ß = .0004, SE = .0002, *t* = 2.44, *p* = .015. The age-IVS relationship was also directionally positive in the left hippocampus but was not significant, ß = .0002, SE = .0002, *t* = 0.90, *p* = .367. There was also a significant negative relationship between age and IVS in the left hippocampal body, ß = -.0003, SE =.0002, *t* = -1.97, *p* = .0499. However, the zero-order correlation in this region did not approach significance (r = -.04, p = .47), meaning that age only predicted IVS once gender, motion, and hippocampal volume were accounted for. No other age-IVS relationship reached significance (p-value range .077, .927). Thus, it seems that the overall age x hippocampal subregion interaction was driven by the most posterior subregion being on a different trajectory than the intermediate and anterior regions, with more correlated signals in older age potentially reflecting dedifferentiation of posterior hippocampal signals.

Next, we sought to relate differences in intra-hippocampal signals to differences in episodic memory performance. To do so, we first computed a base linear regression model with age plus standard covariates as predictors of episodic memory. That model showed a strong and significant age-episodic memory relationship, ß = -.033, *t* = -9.55, *p* < .0001, consistent with the zero-order age-episodic memory correlation of r = -.63 from Figure 2. We then computed a full model that added the intervoxel similarity of each hippocampal subregion and the age x intervoxel similarity interaction effects. To simplify the regression model and because right and left IVS for a given hippocampal subregion were correlated with one another (tail r = .375, body r = .331, head r = .523, all p’s < .001), we averaged IVS values across hemispheres for each subregion. Results from the full model are presented in Table 2. While the overall model was quite successful in accounting for differences in episodic memory, *F*(11,145) = 11.64, *p* < .001, adjusted *R^2^*= .41, none of the individual IVS predictors or age x IVS interaction effects were significant, and there remained a strong negative relationship between age and episodic memory with IVS accounted for. Further, the full model was not a significant improvement over the age only model, *F*(6,145) = .49, *p* = .81. This lack of a significant IVS-episodic memory relationship remained when we ran separate regressions for each hippocampal subregion (all IVS and age x IVS interaction effects p > .24), when we ran separate regressions for the right and left hemispheres (all IVS and age x IVS interaction effects p > .15), and when we removed covariates (all IVS and age x IVS interaction effects p > .24). Thus, despite signs of age-related differences in the similarity of signals within the hippocampus, intra-hippocampal signals at rest were not reliable predictors of age-related differences in episodic memory performance.

**Table 2:**
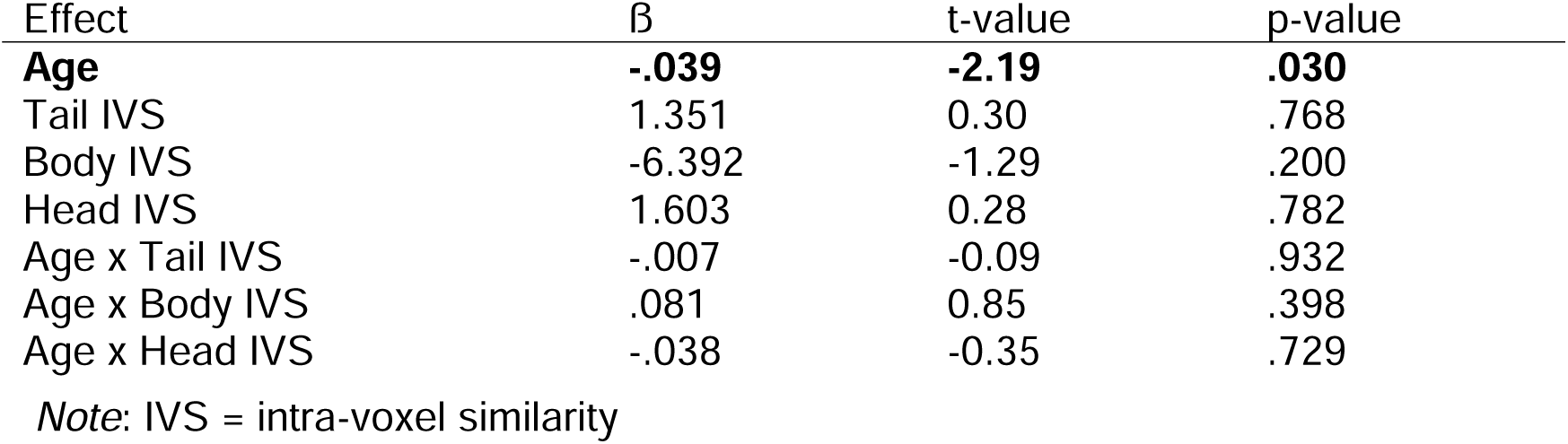
Multiple regression relating intervoxel similarity in each hippocampal region to episodic memory performance.

| Effect | $\beta$ | t-value | p-value |
| --- | --- | --- | --- |
| **Age** | **-.039** | **-2.19** | **.030** |
| Tail IVS | 1.351 | 0.30 | .768 |
| Body IVS | -6.392 | -1.29 | .200 |
| Head IVS | 1.603 | 0.28 | .782 |
| Age x Tail IVS | -.007 | -0.09 | .932 |
| Age x Body IVS | .081 | 0.85 | .398 |
| Age x Head IVS | -.038 | -0.35 | .729 |

*Note:* IVS = intra-voxel similarity

### 3.2. Age-related differences in hippocampal-whole brain functional connectivity

#### 3.2.1 Similarity in connectivity profiles

Looking beyond intra-hippocampal signals, functional specialization along the hippocampal long axis has been shown to emerge in part due to different patterns of connectivity with the rest of the brain (Catenoix et al., 2011; Duvernoy et al., 2005; Fanselow & Dong, 2010; L. E. Frank et al., 2019; Jung et al., 1994; Kahn et al., 2008). To test for age-related differences in patterns of functional connectivity along the hippocampal longitudinal axis, we computed functional connectivity between each hippocampal subregion and 42 cortical and subcortical regions in each hemisphere (84 total targets across hemispheres) defined from the Desikan-Killiany Atlas (Desikan et al., 2006). Figure 4 depicts functional connectivity values for all target regions separated by hippocampal subregion and approximately decade-based age groups (18-29, 30-39, 40-49, etc.). Qualitatively, we saw expected patterns such as stronger ipsilateral connectivity compared to contralateral connectivity, generating the checkerboard pattern within each hippocampal-target ROI pair. We saw that the hippocampal body had the strongest connectivity with the parahippocampal gyrus, whereas the hippocampal head had the strongest connectivity with the amygdala. For both the body and head, the supramarginal gyrus was the strongest negative connection. The hippocampal tail showed more distributed connectivity and did not have a single, clear strongest connection. We also saw that there was not one predominant age-related difference that was consistent and visible across many connections. Instead, the direction of the connections was relatively consistent across age groups for most ROI pairs with subtle differences in their strength.

**Figure 4.**
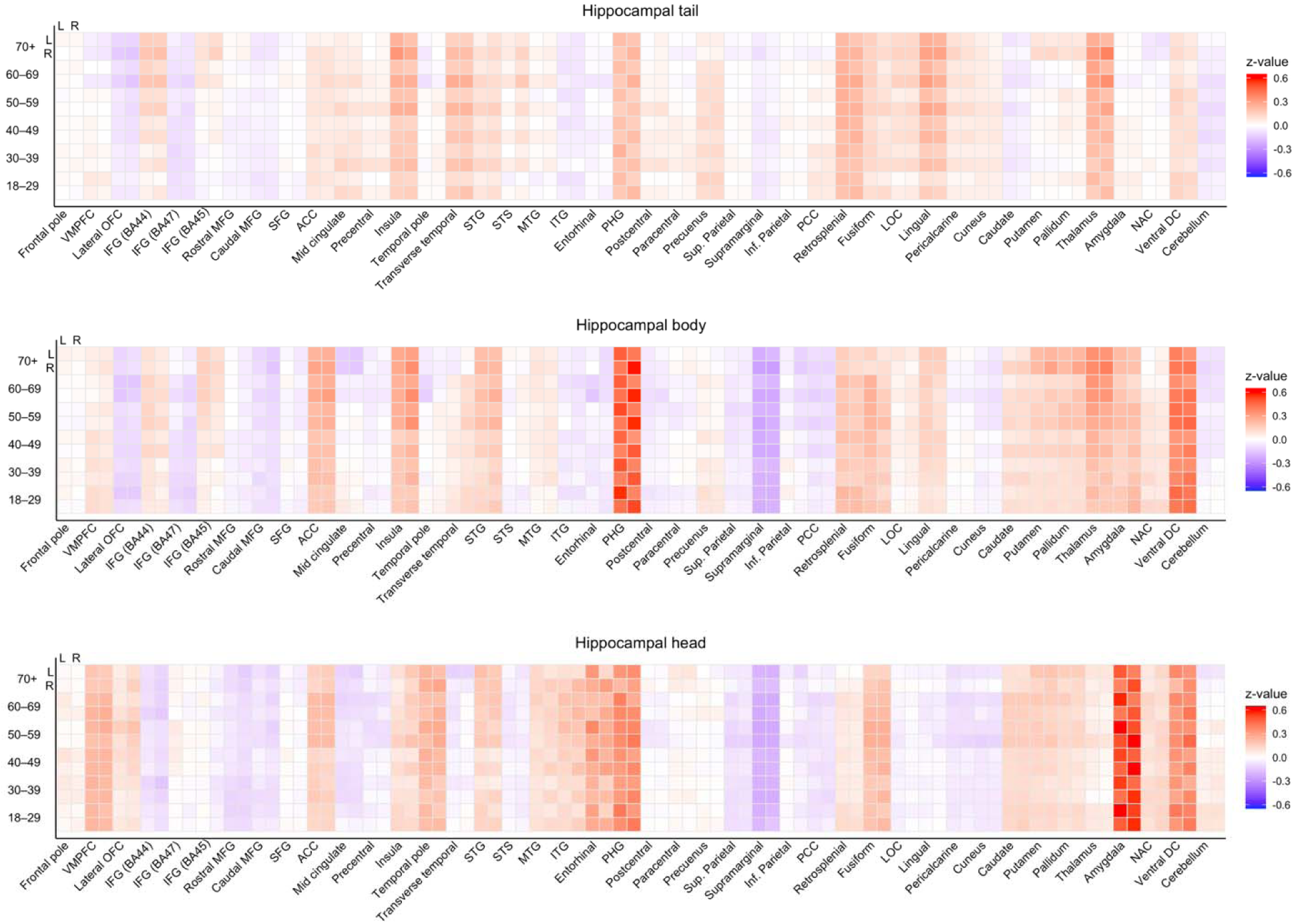
Functional connectivity for each hippocampal subregion and target region across the adult lifespan. Functional connectivity values are presented separately for the hippocampal tail, hippocampal body, and hippocampal head. Functional connectivity was computed as the Fisher-z transformed partial correlation between timecourses in each hippocampal seed region and 84 cortical and subcortical target regions across both hemispheres. Darker red colors represent stronger positive connections, darker blue colors represent stronger negative connections, and white represents connectivity values near zero. Participants were separated into categorical age groups to visualize age effects. Left (L) and right (R) hippocampal seeds are presented in adjacent rows, and left and right target regions are presented in adjacent columns. Full names and atlas labels for abbreviated target regions are presented in Supplementary Table S1.

Our first approach to quantifying patterns of hippocampal connectivity was to determine the degree of overlap in the whole-brain connectivity profiles of each pair of hippocampal subregions and whether similarity in connectivity profiles differed across the adult lifespan. We used an approach similar to representational similarity analyses that are typically used to determine the similarity between patterns of functional activation (Kriegeskorte et al., 2008), substituting patterns of connectivity strength across target regions. In each individual subject, we computed the correlation between the unthresholded pattern of connections for each pair of ipsilateral hippocampal segments (e.g., left tail connectivity map similarity to left body connectivity map). This process resulted in six Fisher’s Z transformed correlation values for each subject (tail-body, body-head, tail-head for left and right hippocampus). To test for differences in the degree of connectivity overlap between subregions of the hippocampus, we submitted these connectivity similarity values to a linear mixed effects model that included the region comparison (tail-body, body-head, tail-head), hemisphere, and age as predictors of interest alongside standard covariates. There was a significant overall effect of the regions being compared, ß = -.258, *t* = -5.48, *p* < .0001. Figure 5A depicts the mean similarity in connectivity profiles for each subregion comparison separately for each hemisphere, collapsed across all ages. All pairwise comparisons between connectivity profile values were significant (all t’s > 3.5, p’s < .001, d’s > 2.6). Overall, the connectivity profile for the body of the hippocampus was most similar to the tail, but the body also showed strong similarity to the connectivity profile of the head. This finding is notable since a common scheme for splitting the hippocampus into two segments collapses across the body and tail of the hippocampus in defining the posterior subregion and leaves the head as the anterior subregion (Poppenk et al., 2013). Our results suggest that the body may be more of a mix of ‘tail-like’ ‘head-like’ connectivity, with a slight bias toward the hippocampal tail. In contrast, the overlap in connectivity profiles was substantially lower for the tail and head, representing the extreme posterior and extreme anterior hippocampal subregions. Together, these findings are in line with different functional properties along the hippocampal long axis being related to their patterns of connectivity with the rest of the brain.

**Figure 5.**
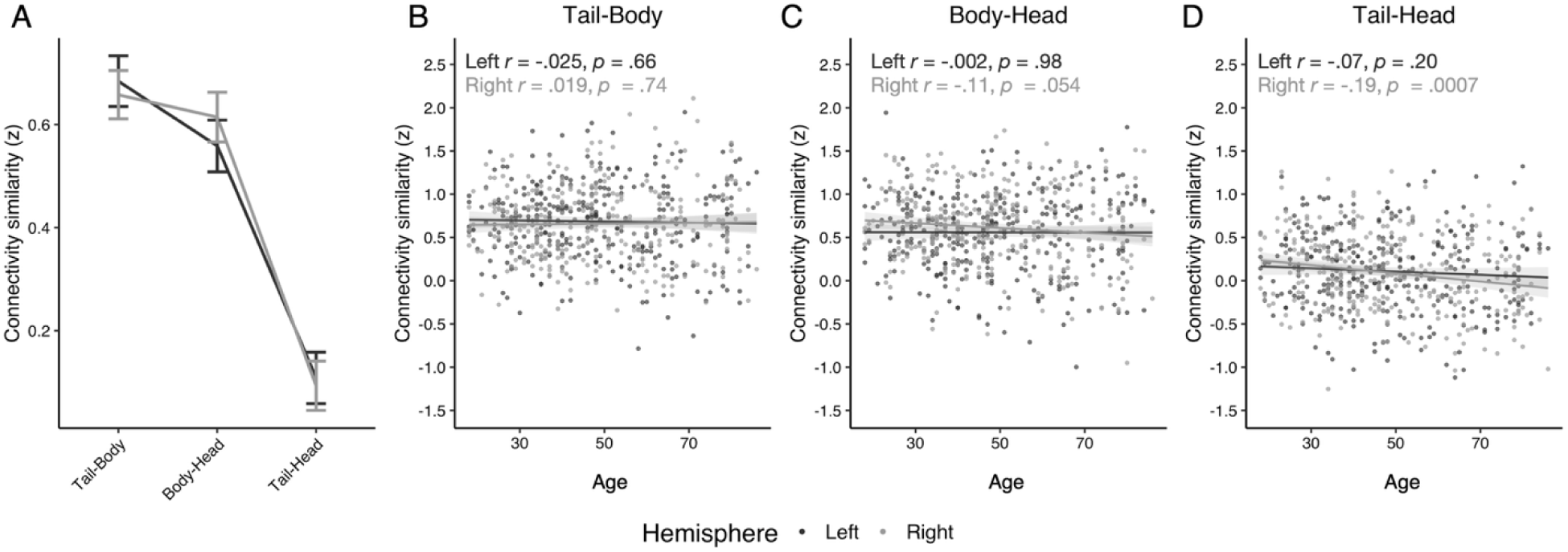
Similarity in connectivity profiles between hippocampal subregions. (A) Similarity in unthresholded functional connectivity profiles between pairs of hippocampal subregions. The relationship between continuous age (in years) and similarity in unthresholded functional connectivity profiles between the (B) tail and body, (C) body and head and (D) tail and head. Similarity values are the Fisher’s Z transformation of the Pearson’s correlation between the connectivity values for all target regions across both hemispheres associated with each pair of hippocampal subregions. Results are presented separate for the left (dark) and right (light) hippocampus. Error bars/shaded represent 95% confidence intervals.

The overall effect of age on the similarity of connectivity profiles was not significant, ß = -.0001, *t* = -.066, *p* = .947, nor was the age x region comparison interaction, ß = -.0006, *t* = -.68, *p* = .498. These findings suggest that differences in connectivity profiles along the hippocampal long axis were relatively preserved in older age, consistent with our qualitative conclusion from Figure 4. Yet, there was a trend toward a three-way age x region comparison x hemisphere effect that did not reach significance, *ß* = -.002, *t* = -1.59, *p* = .111. In order to better understand this trend, we computed the zero-order relationship between age and similarity scores separately for the tail-body (Figure 5B), body-head (Figure 5C), and tail-head comparisons (Figure 5D) separately for the left and right hemispheres. While most age effects were near zero and did not approach significance, the similarity in connectivity profiles for the right tail and head were significantly less similar in older age (an effect that passed a strict Bonferroni correction for the six comparisons, alpha = .008). The age effect in the left hemisphere was also negative but not significant. The right body and head comparison also showed a similar pattern of reduced overlap in older age that did not reach significance. Taken together, there were differences in connectivity profiles across the hippocampal long axis and that the degree of overlap in the connectivity profiles was relatively stable across the adult lifespan. However, there was some evidence that, if anything, connectivity profiles of the tail and head of the hippocampus became *more differentiated* from one another in older age.

We then tested the extent to which the similarity of connectivity profiles between pairs of hippocampal subregions were related to age differences in episodic memory performance. We compared an age only model to predict episodic memory with a full model that additionally included the connectivity similarity values for each pair of hippocampal subregions (averaged across hemispheres) and their interactions with age. The results from the full model are presented in Table 3. The full model was only a marginally better fit than the age only model, *F*(6,145) = 2.11, *p* = .055, but we nonetheless saw signs of age-related effects in the relationship between connectivity profile similarity and episodic memory. There was a significant negative relationship between tail-body similarity and episodic memory, meaning that individuals with stronger overlap in connectivity profiles between the tail and the body showed poorer memory performance. That relationship was also significantly positively moderated by age, meaning that the relationship was *less negative* in older age. When we computed the similarity-episodic memory relationship separately for those aged 18-49 (n = 90) and those aged 50+ (n = 66), there was a marginally significant negative relationship among the younger subsample (*ß* =-.565, *t* = -1.86, *p* = .067) and a numerically positive but non-significant relationship in the older subsample (*ß* = .286, *t* = 1.061, *p* = .293). Thus, having more similar connectivity profiles between the tail and the body of the hippocampus was an indicator of poorer performance in younger age, but that became less true in older age. Yet, we also saw that the age effect in episodic memory was not explained by the similarity between connectivity profiles and actually got somewhat stronger (from -.033 to -.052) once connectivity profiles were considered. This stronger age effect is likely because the tail-body similarity was primarily explaining age differences in memory performance among the younger portion of the sample, leaving the larger performance differences between the young and older portions of the sample unaccounted for.

**Table 3:** Multiple regression relating the similarity connectivity profiles to episodic memory performance.

| Effect | $\beta$ | t-value | p-value |
| --- | --- | --- | --- |
| **Age** | **-.052** | **-5.79** | **< .001** |
| **Tail-body similarity** | **-1.22** | **-2.23** | **.027** |
| Body-head similarity | -.745 | -1.39 | .168 |
| Tail-head similarity | .411 | 0.600 | .552 |
| **Age x Tail-body similarity** | **.023** | **2.328** | **.021** |
| Age x Body-head similarity | .008 | 0.845 | .400 |
| Age x Tail-head similarity | .001 | 0.064 | .949 |

*Note:* bold text represents effects passing an alpha = .05 threshold.

#### 3.2.2 Age differences in the strength of individual hippocampal subregion connections

Next, we sought to understand the specific connections that differed between hippocampal subregions and any age-related differences in the pattern of connectivity across hippocampal subregions and the rest of the brain. For each of the 42 target ROIs, we computed a linear mixed effects model that included hippocampal subregion (tail, body, head), hippocampal hemisphere (left, right), and target ROI hemisphere (left, right) as within-subject fixed effects. A random intercept and random slopes by subject were included for each within-subject variable. We included age as a between-subjects fixed effect. We also included the age x hippocampal subregion interaction in the model because age differences in hippocampal connectivity were of key interest. We did not include other interaction effects in order to simplify the model, but we included standard gender, motion, and hippocampal volume as fixed effect covariates. We then computed F-tests on this model to test for any difference among hippocampal subregions and any age moderation of those patterns. Results from the main effect of hippocampal ROI and the age x hippocampal ROI interaction are presented in Table 4.

**Table 4:**
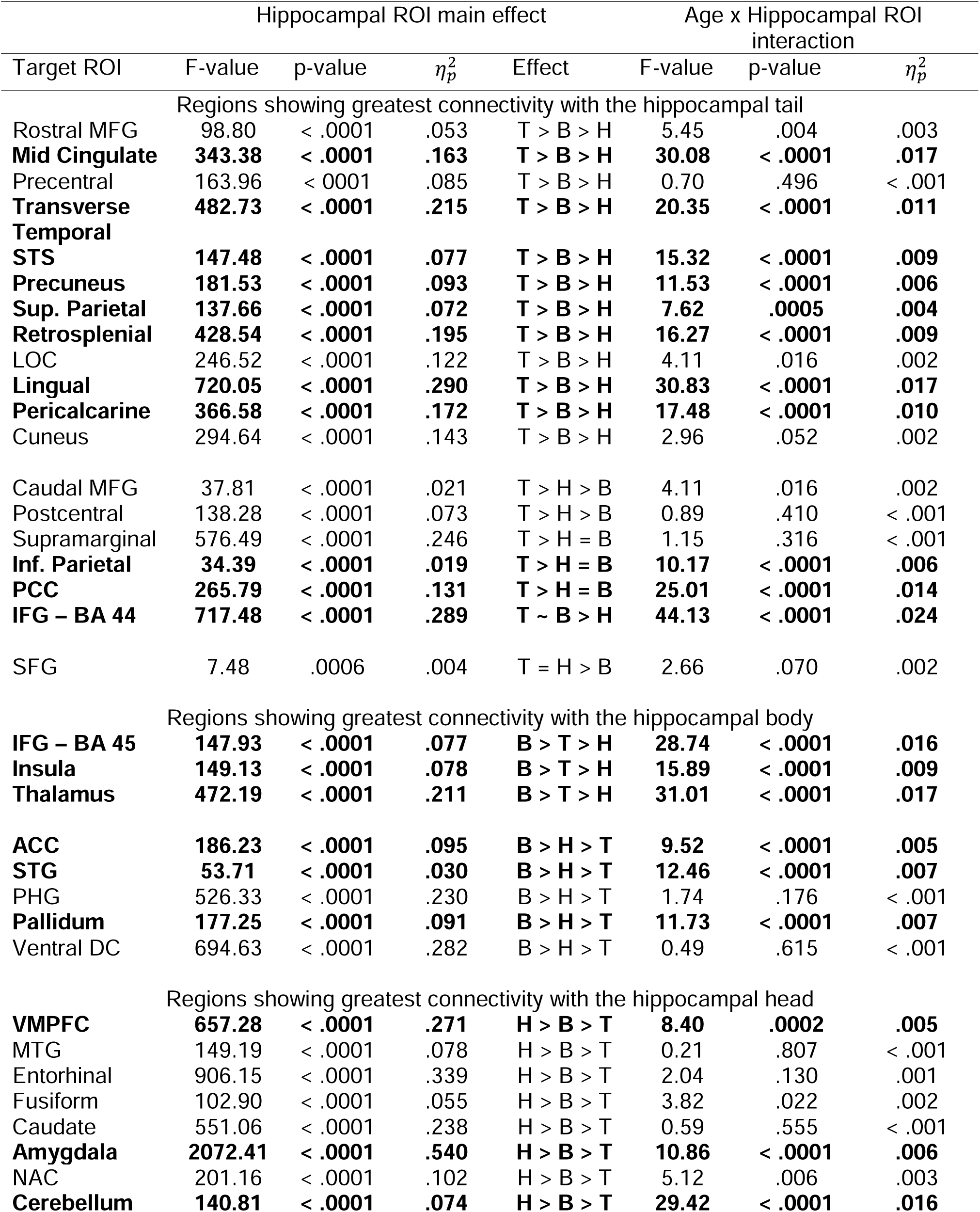
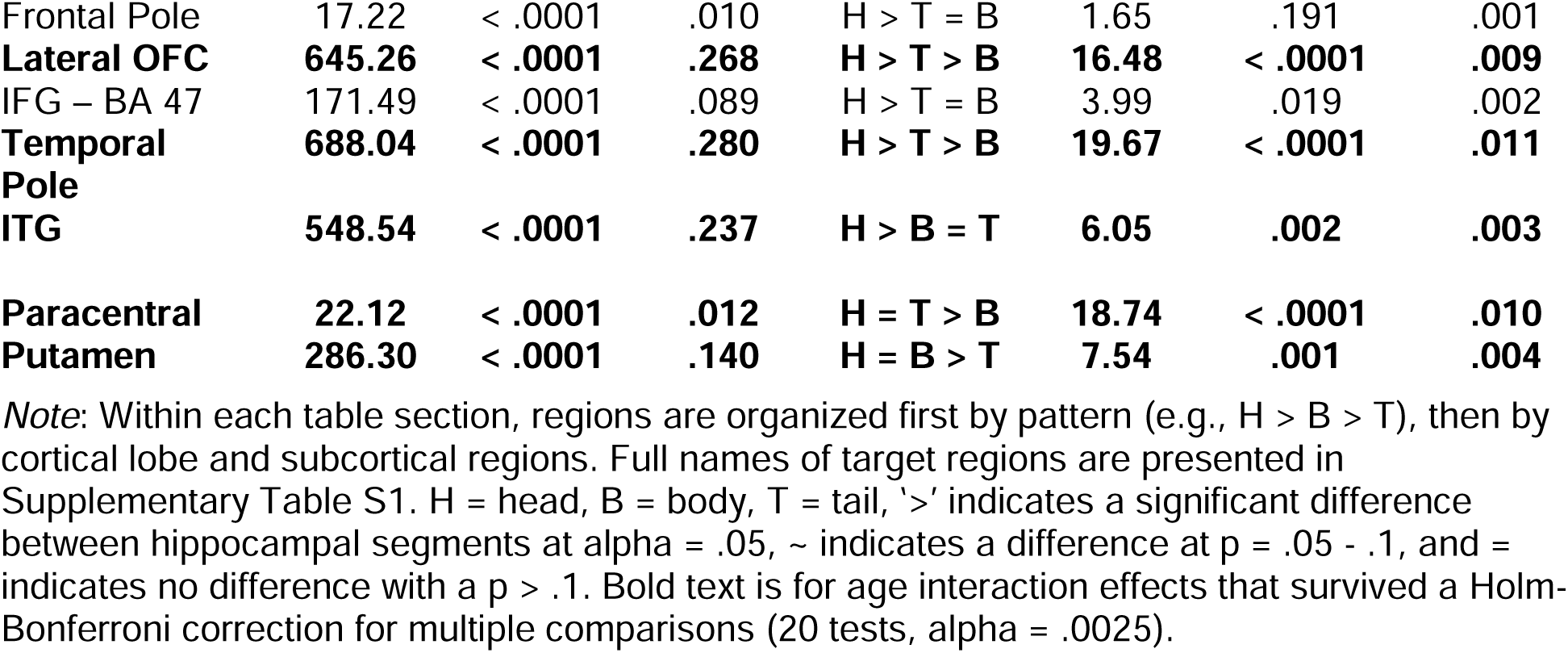
F-tests for overall differences in connectivity along the hippocampal long axis and age moderation of those effects.

All target regions showed connectivity differences across hippocampal subregions that passed a corrected threshold (Bonferroni-Holm correction for 42 tests, alpha = .00119). Thus, differences in the strength of functional connections across the hippocampal long axis were robust. The most common single pattern was a posterior to anterior gradient of decreasing connectivity (i.e., tail > body > head). Twelve regions (29% of regions) showed that pattern, which included a number of visual and parietal regions. An additional 7 regions (17% of regions) showed connectivity that was at least numerically strongest for the hippocampal tail but with a different specific pattern. The next most common pattern was the reverse gradient: increasing connectivity from posterior to anterior (8 regions, 19% of regions). This set of regions included the VMPFC, middle temporal gyrus, entorhinal cortex, fusiform cortex, and the amygdala. An additional 7 regions (17% of regions) showed connectivity that was at least numerically strongest for the hippocampal head but with a different specific pattern. Lastly, a total of 8 regions (19% of regions) showed the strongest connectivity with the hippocampal body, with 3 regions (7% of regions) showing their next strongest connectivity with the hippocampal tail and the other 5 regions (12% of regions) showing their next strongest connectivity with the hippocampal head. Thus, across the 42 regions tested, there were clear differences in the strength of connections to subregions of the hippocampus, clear posterior-to-anterior gradients, and a smaller subset of regions showing the strongest connectivity with the hippocampal body.

Next, we followed-up on the relationship between age and connectivity strength differences across hippocampal subregions for each of the target regions where the age x hippocampal subregion interaction effect passed a correction for multiple comparisons (as noted in Table 4). The partial correlations between age and connectivity strength between each target ROI and each hippocampal subregion accounting for subject-level motion are presented in Table 5. Because these were post-hoc tests to probe significant interactions, we did not impose a multiple comparisons correction.

**Table 5:**
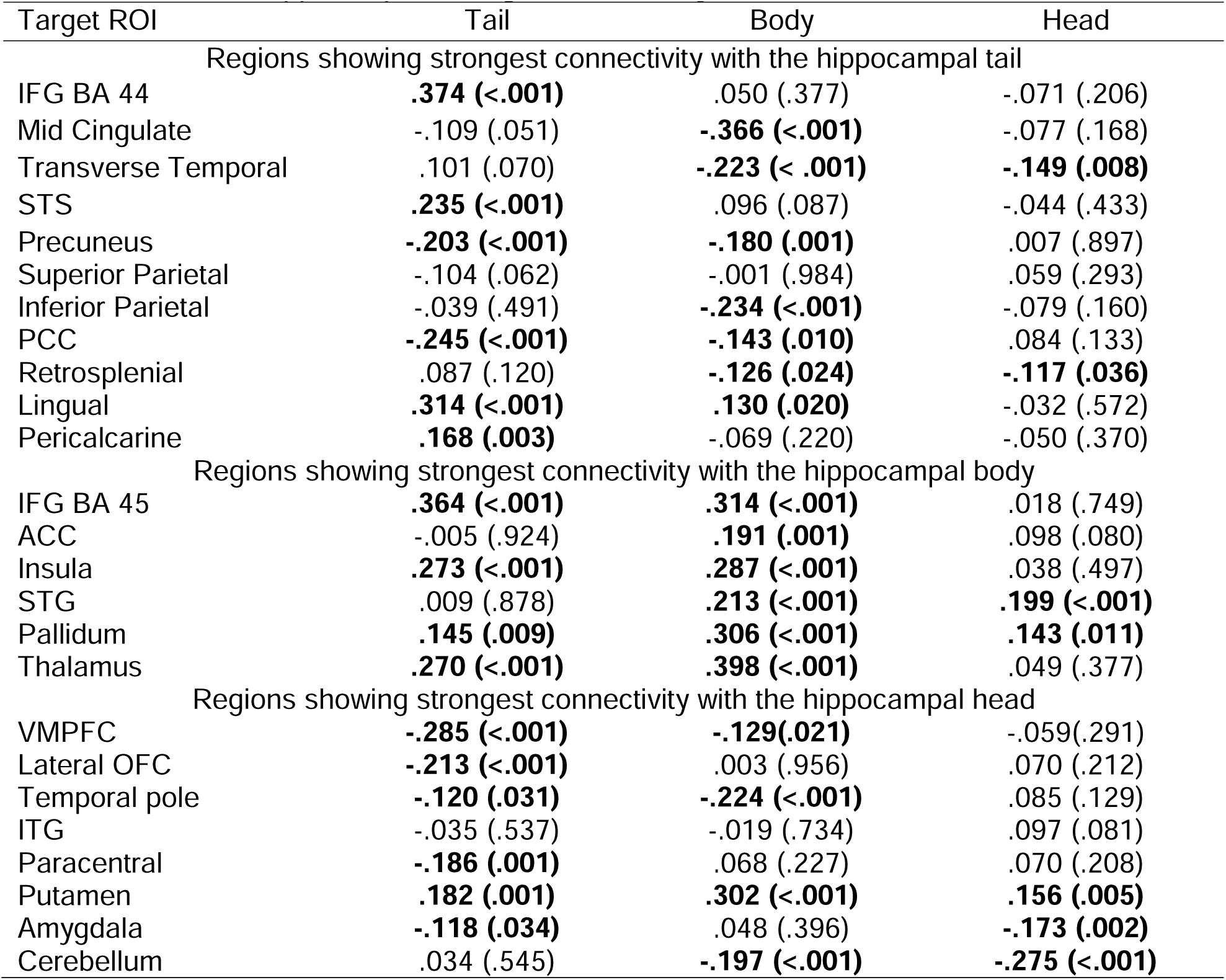
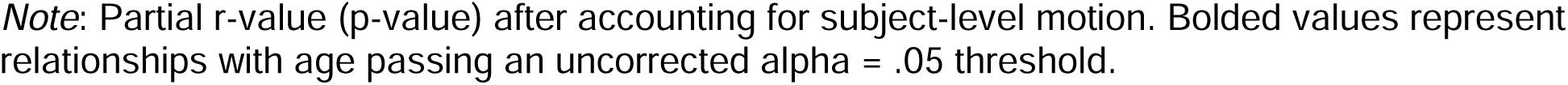
Relationship between age and connectivity strength among target ROIs showing age moderation of their hippocampal subregion connectivity.

| Target ROI | Tail | Body | Head |
| --- | --- | --- | --- |
| Regions showing strongest connectivity with the hippocampal tail |  |  |  |
| IFG BA 44 | **.374 (<.001)** | .050 (.377) | -.071 (.206) |
| Mid Cingulate | -.109 (.051) | **-.366 (<.001)** | -.077 (.168) |
| Transverse Temporal | .101 (.070) | **-.223 (<.001)** | **-.149 (.008)** |
| STS | **.235 (<.001)** | .096 (.087) | -.044 (.433) |
| Precuneus | **-.203 (<.001)** | **-.180 (.001)** | .007 (.897) |
| Superior Parietal | -.104 (.062) | -.001 (.984) | .059 (.293) |
| Inferior Parietal | -.039 (.491) | **-.234 (<.001)** | -.079 (.160) |
| PCC | **-.245 (<.001)** | **-.143 (.010)** | .084 (.133) |
| Retrosplenial | .087 (.120) | **-.126 (.024)** | **-.117 (.036)** |
| Lingual | **.314 (<.001)** | **.130 (.020)** | -.032 (.572) |
| Pericalcarine | **.168 (.003)** | -.069 (.220) | -.050 (.370) |
| Regions showing strongest connectivity with the hippocampal body |  |  |  |
| IFG BA 45 | **.364 (<.001)** | **.314 (<.001)** | .018 (.749) |
| ACC | -.005 (.924) | **.191 (.001)** | .098 (.080) |
| Insula | **.273 (<.001)** | **.287 (<.001)** | .038 (.497) |
| STG | .009 (.878) | **.213 (<.001)** | **.199 (<.001)** |
| Pallidum | **.145 (.009)** | **.306 (<.001)** | **.143 (.011)** |
| Thalamus | **.270 (<.001)** | **.398 (<.001)** | .049 (.377) |
| Regions showing strongest connectivity with the hippocampal head |  |  |  |
| VMPFC | **-.285 (<.001)** | **-.129(.021)** | -.059(.291) |
| Lateral OFC | **-.213 (<.001)** | .003 (.956) | .070 (.212) |
| Temporal pole | **-.120 (.031)** | **-.224 (<.001)** | .085 (.129) |
| ITG | -.035 (.537) | -.019 (.734) | .097 (.081) |
| Paracentral | **-.186 (.001)** | .068 (.227) | .070 (.208) |
| Putamen | **.182 (.001)** | **.302 (<.001)** | **.156 (.005)** |
| Amygdala | **-.118 (.034)** | .048 (.396) | **-.173 (.002)** |
| Cerebellum | .034 (.545) | **-.197 (<.001)** | **-.275 (<.001)** |

*Note:* Partial r-value (p-value) after accounting for subject-level motion. Bolded values represent relationships with age passing an uncorrected alpha = .05 threshold.

There were 11 regions that showed their strongest connectivity with the hippocampal tail that also showed age-related moderation of hippocampal connectivity. The relationship between age and connectivity strength to each hippocampal subregion for each of these regions is depicted in Figure 6. A common pattern across regions was an overall differentiation such that connectivity differences between the hippocampal subregions were exaggerated in older age. Seven regions, IFG BA 44, transverse temporal cortex, STS, inferior parietal, retrosplenial cortex, lingual gyrus, and pericalcarine cortex, showed that overall pattern. Two regions, the precuneus and superior parietal cortex showed the opposite pattern: a convergence of connectivity strength in older age such that differences between hippocampal subregions were reduced in older age. Across target regions, there was a lack of significant age-related differences in connectivity with the hippocampal head: only two regions (transverse temporal cortex and retrosplenial cortex) showed significant age-related differences in the strength of connectivity with the hippocampal head (decreases in both cases), whereas seven regions showed age-related connectivity differences with the body, and six regions showed differences with the tail. Thus, among these target regions, hippocampal head connectivity was less impacted by older age than hippocampal body and tail connectivity.

**Figure 6.**
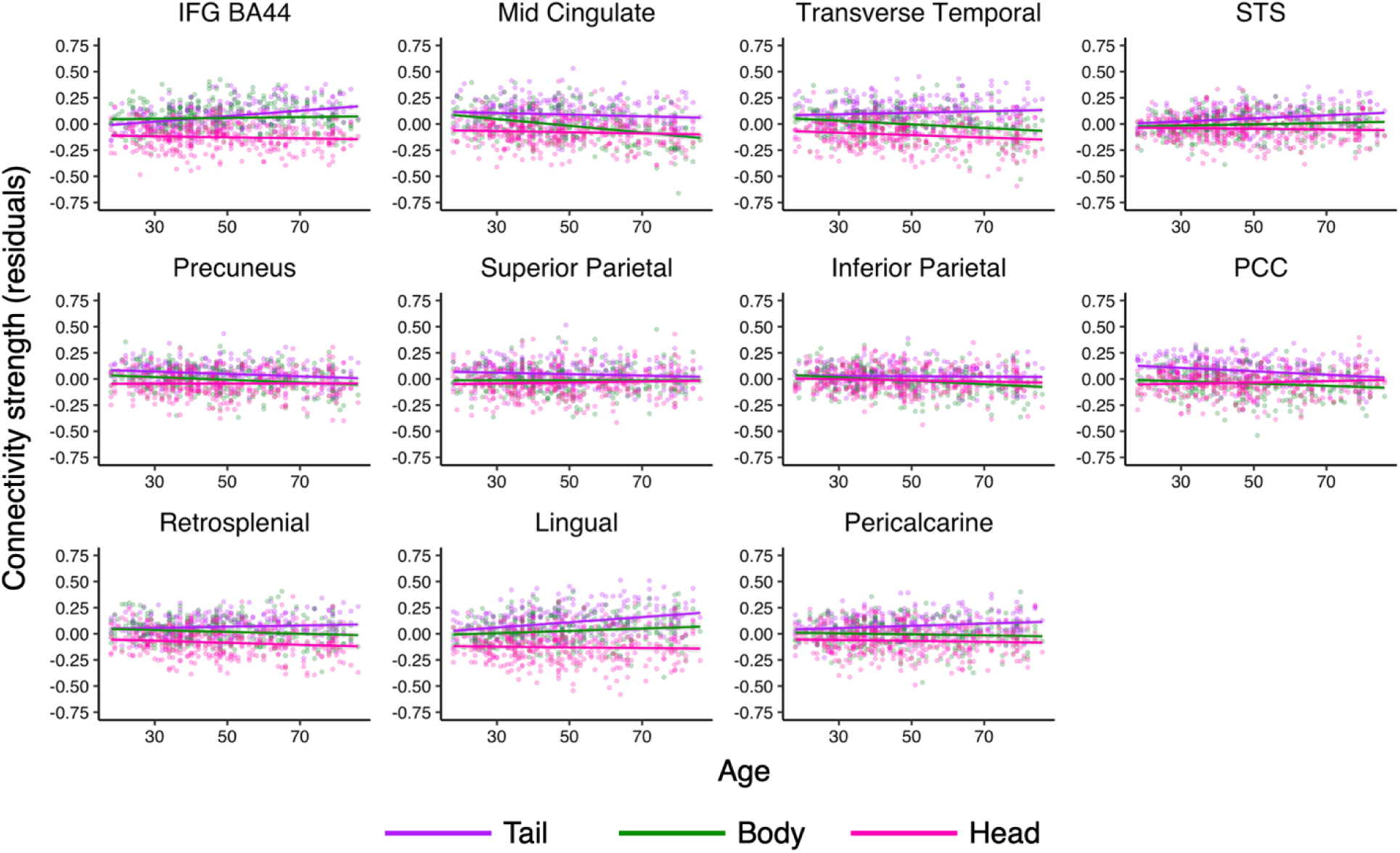
Age effects in functional connectivity with hippocampal subregions in regions most connected to the hippocampal tail. The relationship between continuous age (in years) and functional connectivity strength for each hippocampal subregion and the target regions that showed strongest connectivity with the hippocampal tail and significant age moderation. Dots represent individual subjects and lines represent linear age trends. The y-axis is connectivity strength after accounting for subject-wise motion (i.e., residuals from a linear model relating connectivity strength and motion).

There were six regions that had the strongest connectivity with the hippocampal body and also showed an age interaction. The relationship between age and connectivity strength to each hippocampal subregion for each of these regions is depicted in Figure 7. There was a very consistent age-related pattern in regions most connected to the body: significant age-related increases in connectivity with the hippocampal body. All but one region (STG) also showed significant age-related increases in connectivity with the hippocampal tail, and two regions (STG and pallidum) showed increased connectivity with the hippocampal head. None of these regions showed significant age-related decreases in connectivity strength. The net result was increased connectivity differentiation among hippocampal subregions later in life. While the age-related pattern is clear, the target regions showing this pattern do not have a clear functional theme among them, with the pallidum typically linked motor function (Lanciego et al., 2012), the thalamus containing multiple nuclei involved in different sensory, motor and cognitive functions (Fama & Sullivan, 2015), the insula typically associated with attentional salience (Menon & Uddin, 2010), and STG and IFG (BA 45) involved in language processing (Stefaniak et al., 2021). Thus, we see the body of the hippocampus having increased connectivity with a heterogeneous set of cortical and subcortical regions in older age.

**Figure 7.**
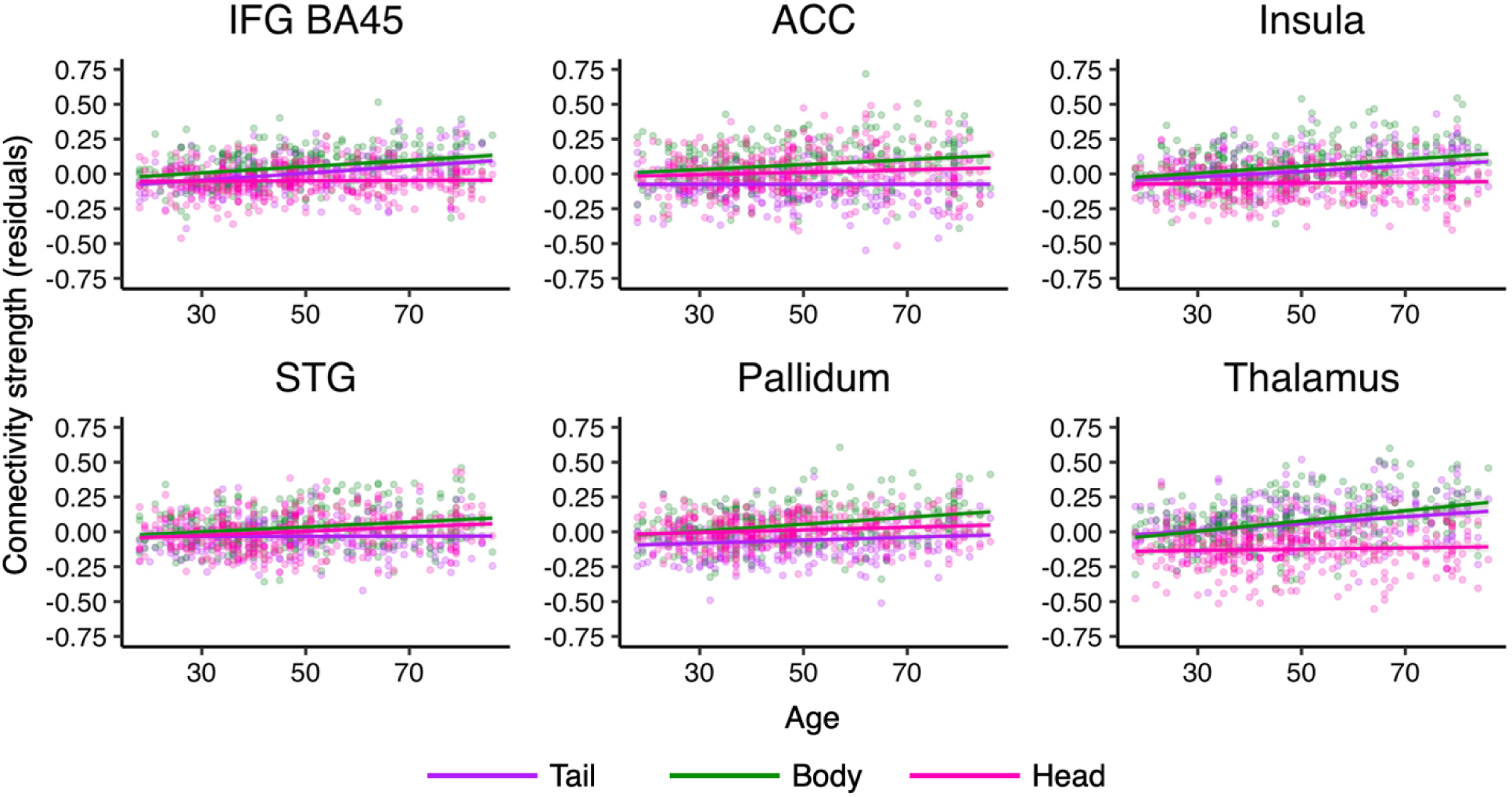
Age effects in functional connectivity with hippocampal subregions in regions most connected to the hippocampal body. The relationship between continuous age and functional connectivity strength for each hippocampal subregion and the target regions that showed strongest connectivity with the hippocampal body and significant age moderation. Dots represent individual subjects and lines represent linear age trends. The y-axis is connectivity strength after accounting for subject-wise motion (i.e., residuals from a linear model relating connectivity strength and motion).

There were eight regions that had the strongest connectivity with the hippocampal head and also showed an age interaction. The relationship between age and connectivity strength to each hippocampal subregion for each of these regions is depicted in Figure 8. In these regions, there was also a fairly consistent pattern: age-related reductions in connectivity strength, most commonly with the hippocampal tail (five regions), but also with the body (three regions) and head (two regions). Only the putamen showed significant age-related increases in connectivity and did so across all three subregions (but most strongly with the body). However, the *net effect* of age-related differences across the hippocampal subregions was not consistent across regions. The VMPFC and, to some extent, the temporal pole showed increasing differentiation of hippocampal connectivity with increasing age. The cerebellum showed decreasing differentiation of connectivity with increasing age. All other regions showed a pattern where one or two of the hippocampal subregions were on a different age trajectory than the remaining subregion(s), but the exact pattern differed across targets. Yet, consistent with the pattern from regions most connected to the hippocampal tail and body, we see the fewest age-related differences in the strength of connectivity to the hippocampal head, suggesting connectivity with this hippocampal subregion is less impacted in older age regardless of its overall level of connectivity compared to other subregions.

**Figure 8.**
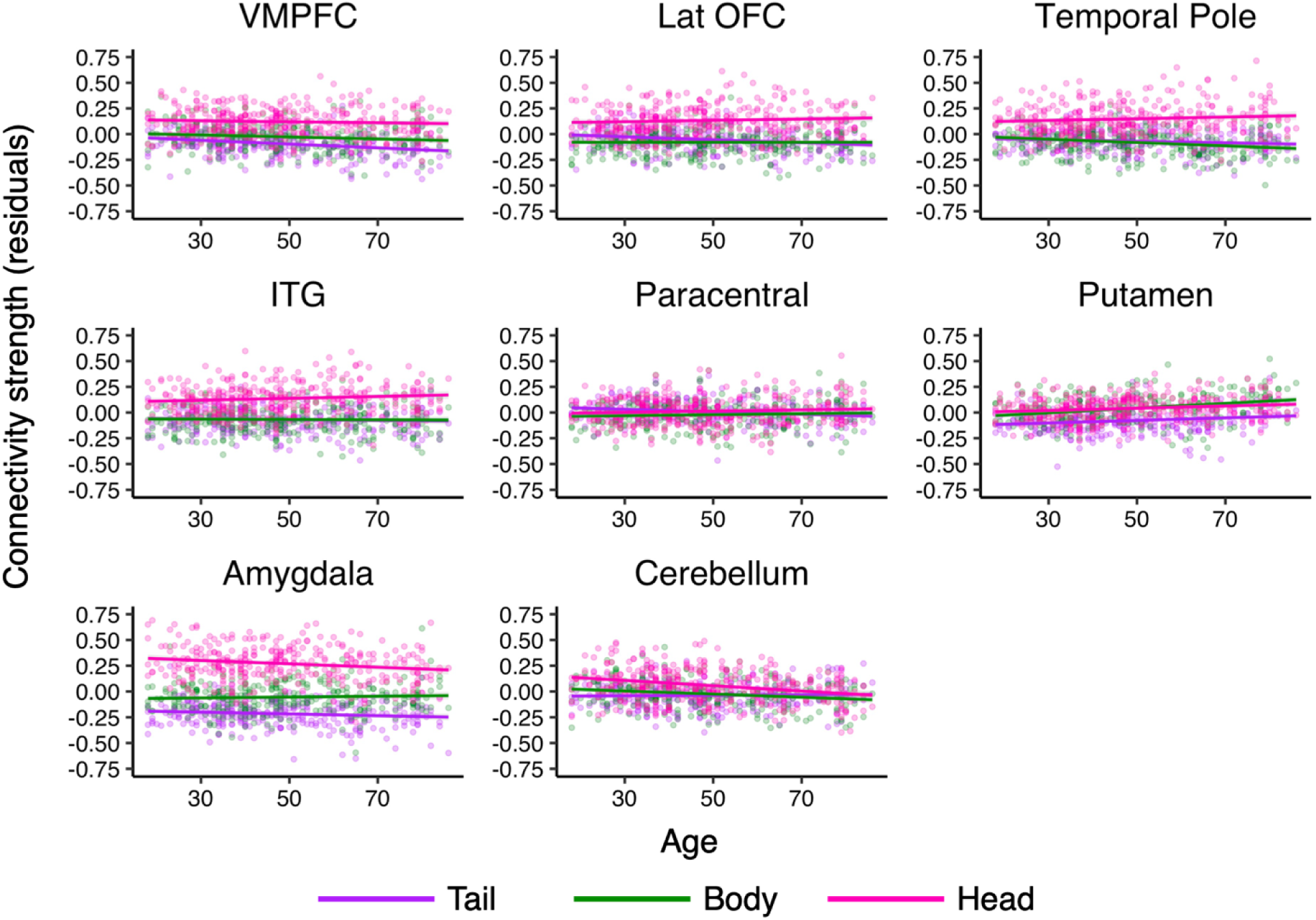
Age effects in functional connectivity with hippocampal subregions in regions most connected to the hippocampal head. The relationship between continuous age (in years) and functional connectivity strength for each hippocampal subregion and the target regions that showed strongest connectivity with the hippocampal head and significant age moderation. Dots represent individual subjects and lines represent linear age trends. The y-axis is connectivity strength after accounting for subject-wise motion (i.e., residuals from a linear model relating connectivity strength and motion).

Lastly, we tested the extent to which hippocampal connectivity could explain age differences in episodic memory performance. Our first approach was to test whether any individual connections served to mediate the relationship between age and episodic memory scores. We computed separate models that added connectivity between one of the hippocampal subregions and one of the target regions to a base age model, testing whether the age effect in episodic memory would remain with each connection considered. Unsurprisingly given the complexity of episodic memory and robustness of its age-related decline, no single connection accounted for the age difference nor made any meaningful reduction in the age effect on episodic memory (all age ß’s < -.308, *p*’s < .0001). Figure 9A depicts the zero order correlations between the individual hippocampal – target ROI connections and episodic memory scores. Here, the issue becomes clear: while the zero order correlation between episodic memory and age was strong (r = -.63), the correlations between connectivity and episodic memory were relatively weak, ranging from r = -.24 (Tail-Fusiform) to .27 (Tail-Ventral DC).

**Figure 9.**
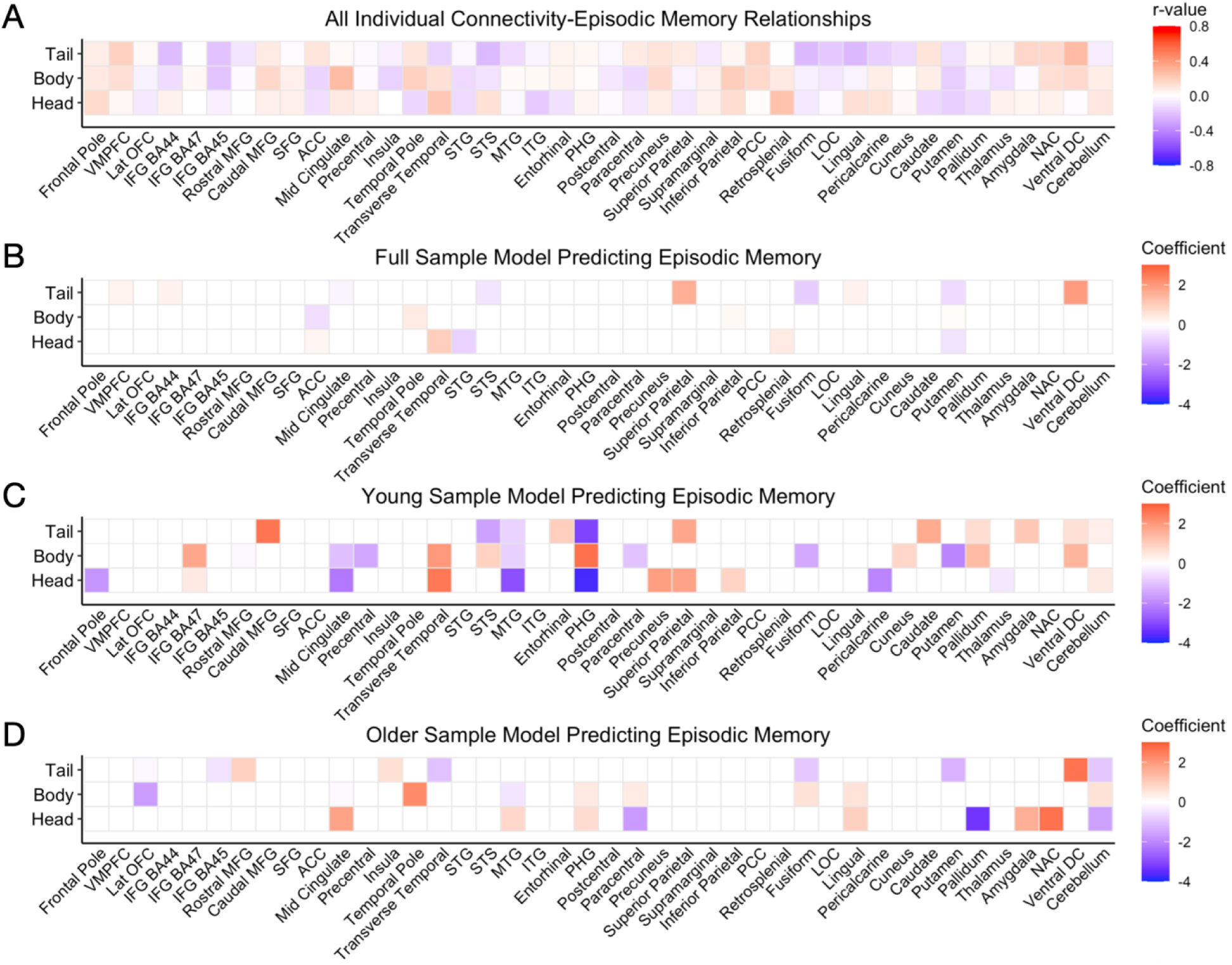
Relationships between episodic memory performance and connectivity strength. (A) The zero order relationship between episodic memory performance and functional connectivity strength for each hippocampal subregion-target region combination. (B) Coefficients from a model predicting episodic memory scores from hippocampal functional connectivity across the full sample. (C) Coefficients from a model predicting episodic memory scores from hippocampal functional connectivity in the younger half of the sample (aged 18-49, n = 90). (D) Coefficients from a model predicting episodic memory scores from hippocampal functional connectivity in the older half of the sample (aged 50+, n = 66). In A-D, darker red colors represent stronger positive relationships, darker blue colors represent stronger negative relationships, and white represents relationship values near zero (i.e., connections not included in a given model for B-D).

Thus, age alone was a much stronger predictor of episodic memory performance than any individual hippocampal connection.

Yet, the possibility remained that a *combination* of connections could explain episodic memory performance and age differences therein. To test that possibility, we conducted an exploratory regression analysis to identify a set of connections that could predict episodic memory performance. Because of the large number of potential predictors (3 hippocampal subregions x 42 target ROIs = 126), we used an elastic net approach to select predictors for the model (using the glmnet package in R; Friedman et al., 2010; Tay et al., 2023). Elastic net regression (Zou & Hastie, 2005) is a compromise between ridge regression, which can handle multicollinearity of predictors well, but retains all predictors in the final model (Hoerl & Kennard, 1970), and lasso regression, which allows coefficients to be set to zero to allow for selection of relevant predictors (Tibshirani, 1996). Using this approach, we first estimated the best fitting model across the entire sample without considering age. Figure 9B depicts the coefficients from the resulting model and Supplementary Table S2 presents the coefficients along with their p-values. We entered the non-zero predictors from this elastic net model into a standard linear model alongside age and compared this age + connectivity model to the base model with only age. The result was a significant improvement in fit, *F*(18,133) = 2.44, *p* = .0019. Thus, age and connectivity together explained differences in episodic memory performance better than age alone. Yet, including functional connectivity did not entirely account for age differences in episodic memory: age continued to be a significant predictor of episodic memory in the age + connectivity model, ß = -.026, *t* = -6.71, *p* < .0001. Nonetheless, there was a 21.5% reduction in the magnitude of the negative age effect (i.e., the coefficient was closer to zero) in the model that considered both age and functional connectivity.

The sparseness of this full model (Figure 9B) led us to test whether young and older adults might rely on different sets of connections to support episodic memory performance, leading to few connections that would predict episodic memory when those groups were combined. To test this idea, we generated separate connectivity-episodic memory elastic net models for the young (aged 18-49, n = 90; Figure 9C) and older subsamples (aged 50+, n = 66; Figure 9D). The full list of included connections and their coefficients and p-values are presented in Supplementary Table S2. First, many more connections were included in the model when it was developed from only the younger sample (38 connections) or only from the older sample (27 connections) compared to the full sample (18 connections). Nonetheless, there was significantly more overlap between the full sample connections and the young adult sample connections than would be expected by chance (dice coefficient = .25, permutation test p < .001). There was also significant overlap between the connections in the full model and the older adult model (dice coefficient = .22, permutation test p < .001), and when we compared the models developed from young and older adults to one another (n connections = 11, dice coefficient = .34, p < .001). Yet, when we considered the *direction* of the relationships among the “overlapping” connections in the young and older models, there was a numerically negative relationship, *r*(9) = -.21, *p* = .54. The largest differences (older ß -younger ß) were for the head-PHG (ß difference = 3.33), head-mid cingulate (ß difference = 2.08), tail-ventral DC (ß difference = 1.88), and head-MTG (ß difference = 1.85). All other differences were < 1 in terms of absolute value. For the three connections with the hippocampal head, the relationship to episodic memory performance was negative in the younger sample and either numerically (PHG, MTG) or significantly (mid cingulate) positive in the older sample. Thus, we see that those in the younger sample who showed stronger connectivity with the hippocampal head tended to perform poorly, whereas that relationship was either no longer present or reversed in the older sample. The relationship between episodic memory and tail-ventral DC connectivity was numerically positive in both samples but significant only in the older sample, suggesting some degree of compensation from the hippocampal tail-ventral DC connection in the older sample to support episodic memory performance.

## 4. Discussion

We aimed to test the degree of functional differentiation along the hippocampal long axis across the adult lifespan and its relationship to episodic memory. We found that signals within the most posterior hippocampal subregion tended to become more similar to one another with increasing age, whereas the intermediate and most anterior regions tended to show less similar intra-region signals in older age. Yet, age-related differences in the similarity of signals within the hippocampus did not significantly relate to episodic memory performance. We found that the whole-brain connectivity profile of the right hippocampal head tended to become less similar to the profiles of the body and tail, whereas the similarity of the body and tail connectivity profiles did not differ with age. Yet, higher similarity between the connectivity profiles of the hippocampal body and tail was associated with poorer episodic memory performance, but only within the younger portion of the sample. When examining individual connections, we found robust connectivity differences along the hippocampal long axis in all target regions tested and significant age-related moderation of hippocampal connectivity in about half of the regions tested. Of the hippocampal subregions, the hippocampal head showed the fewest age-related differences in connectivity strength overall. The net effect of older age on connectivity with the three hippocampal subregions differed by target region, but age-related increases in connectivity differentiation were common, with age-related decreases in connectivity differentiation also present but less common. Finally, we identified several connections with the hippocampal head (PHG, MTG, mid cingulate) and one with the hippocampal tail (ventral DC) that were more positively related to episodic memory performance in older age.

One theory of hippocampal function is that there is a posterior to anterior gradient in which the posterior hippocampal representations are more detailed than anterior hippocampal representations (Poppenk et al., 2013). Compelling evidence for this theory came from a study examining the correlation of signals within the hippocampus, showing less correlated signals in the posterior versus anterior hippocampus (Brunec et al., 2018). The authors concluded that the low correlation among posterior hippocampal signals made the posterior hippocampus well suited to representing information with high resolution to support memory specificity. However, a more recent study tested multiple schemes for subdividing the hippocampus and compared them across two large datasets (Thorp et al., 2022). While there were differences based on the parcellation scheme, results generally pointed to higher similarity among signals in the posterior hippocampus once the hippocampus was subdivided into more than two subregions. Our overall findings are most in line with the latter study: signals within the hippocampal tail were more correlated with one another than signals in the body and the head. One potential driver of differences across studies is that the Brunec et al. (2018) study used both resting state data and data from a navigation task, and the increase in intervoxel similarity from posterior to anterior was more prominent in the navigation task. Our study and the Thorp et al. (2022) study used only resting state data. Thus, it may be that intra-hippocampal signals are noisier during rest and will show more variable patterns across the hippocampal long axis, but that these signaling patterns emerge when the participant is engaged in a task.

Despite an overall pattern of intra-hippocampal signals that differed from predictions, we found that intra-hippocampal signaling in the posterior hippocampus had a different trajectory across the adult lifespan compared to intermediate and anterior regions of the hippocampus, which is consistent with what we would predict based on known age-related declines in various measures of memory specificity (Hashtroudi et al., 1989; Naveh-Benjamin et al., 2003; Yassa et al., 2011). Signals within the posterior hippocampus tended to become more similar to one another with increasing age, the opposite of the trend for the intermediate and anterior hippocampal subregions. This finding is consistent with a prior study showing age-related increases in the strength of functional connectivity within posterior medial temporal lobes (Salami et al., 2016). Increasing similarity of signals in the posterior hippocampus is in line with neurocognitive aging theories suggesting dedifferentiation of neural signals in older age (Goh, 2011; Koen & Rugg, 2019; Park et al., 2012; Pauley et al., 2024): that neural responses become less selective and more broadly tuned in older age. Here we show that this effect may be specific to the posterior versus intermediate and anterior regions of the hippocampus. Following from theories of hippocampal functional specialization, it would make sense for dedifferentiation of posterior hippocampal signals to hinder memory for episodic details. However, we did not find any significant relationship between intra-hippocampal signals and episodic memory performance, nor any age modulation of this relationship. It is possible that the measure of episodic memory did not sufficiently index memory for the kinds of episodic details affected by dedifferentiation of posterior hippocampal signals, or that dedifferentiation of these signals during rest is not predictive of functional specialization during a memory task. However, between the mixed findings on the nature of within-hippocampus signals across its long axis and the lack of a strong behavioral impact of age differences in intra-hippocampal signals, it is also possible that differences in inter-voxel similarity are more an epiphenomenon than a key driver of episodic memory performance.

Yet, evidence for a functional specialization along the hippocampal long axis has also come from differences in patterns of structural and functional connectivity with the rest of the brain. Prior work has shown that the posterior hippocampus has structural and/or functional connections to regions including retrosplenial cortex, the posterior cingulate, precuneus, angular gyrus, visual cortices, and lateral prefrontal cortex (L. E. Frank et al., 2019; Poppenk et al., 2013; Poppenk & Moscovitch, 2011; Ranganath & Ritchey, 2012). In contrast, the anterior hippocampus has shown stronger connectivity with regions including the ventromedial prefrontal cortex, amygdala, and temporal pole (Blessing et al., 2016; Bowman & Zeithamova, 2018; Duvernoy et al., 2013; L. E. Frank et al., 2019; Kier et al., 2004; Robinson et al., 2016, 2016; Wang et al., 2016; Zeithamova et al., 2012). Our results showed robust differences in functional connectivity along the hippocampal long axis that were broadly consistent with the patterns identified in prior work from young adults. We also showed relatively little overlap in connectivity profiles between the most posterior and most anterior hippocampal subregions, whereas the intermediate hippocampal subregion showed considerable overlap with both posterior and anterior hippocampus. Thus, we find strong evidence for a long axis gradient in hippocampal functional connectivity across the lifespan.

We also found that older age moderated the pattern of connectivity across hippocampal subregions for many target regions. Age was most consistently related to connectivity for the hippocampal tail and body, whereas connections with the hippocampal head were less likely to show significant age-related differences. These findings are in line with the hypothesis that relatively posterior hippocampal regions are more impacted by aging (Bussy et al., 2021; Frisoni et al., 2008; Y et al., 2023), but they are novel in that past work on age differences in posterior vs. anterior hippocampal connectivity have been mixed. One prior study used an extreme age groups design and showed age-related reductions in hippocampal functional connectivity with other regions of the medial temporal lobes during a mnemonic similarity task (Stark et al., 2021). Age differences tended to be comparable along the hippocampal long axis (perirhinal cortex) or greater in the anterior versus posterior hippocampus (parahippocampal cortex). Other work also using extreme age groups investigated differences in functional connectivity across the posterior-anterior axis with the rest of the brain and found a shift toward posterior connectivity in older age (Blum et al., 2014). Yet, there were also several regions with age reductions in connectivity with intermediate hippocampal subregions (Blum et al., 2014). Thus, while further empirical evidence is needed to understand the nature of hippocampal long axis connectivity differences in older age, we provide novel evidence that age effects are most prominent in the intermediate and posterior hippocampus and more stable in the anterior hippocampus.

Lastly, we found relatively little consistency in the connections that predicted episodic memory performance across the entire sample and evidence that the direction of the connectivity-performance relationship differed between the younger and older portions of the sample. Three of the connections showing the strongest age differences were with the hippocampal head (PHG, MTG, mid cingulate) while one was with the hippocampal tail (ventral DC). In two cases (PHG, MTG), stronger hippocampal head connectivity was negatively associated with episodic memory performance in the younger subsample, but the relationship was numerically positive in the older subsample. The pattern was similar for the hippocampal head-mid cingulate connection except that the positive relationship in the older subsample was significant. These patterns may have emerged because upregulation of these connections in younger or middle age is essentially showing an ‘older adult’ pattern of connectivity that is maladaptive at that stage in life. Yet, having that pattern in older age was not maladaptive and sometimes even compensatory. We also saw signs of age-related compensation in the connection between the hippocampal tail connection and the ventral DC, an ROI that contains the hypothalamus and subthalamic nucleus (Fischl et al., 2002). Age-related differences in the tail-ventral DC connection may be related to these regions’ involvement in neuromodulatory networks (M. J. Frank, 2006) and neuroendocrine function (Kim & Diamond, 2002) that have key roles in cognition. Taken together, while upregulation of brain activity or connectivity can be a sign of age-related dysfunction (Cabeza et al., 2018; McDonough et al., 2022; Salami et al., 2014), here we see relatively positive links to cognitive performance in the older portion of our sample. However, it is still possible that increasing connectivity particularly to more anterior hippocampal regions had negative consequences for memory performance that were not identified by the episodic memory task used here. Future studies using different memory tasks that tap into a wider range of memory abilities will clarify the implications of maintaining episodic memory performance through upregulation of connectivity with the anterior hippocampus.

## Acknowledgements

This work was supported by internal funding from the University of Wisconsin-Milwaukee, including an Advanced Opportunity Fellowship awarded to C.I.C. The authors thank the members of the Cognition, Aging, and Brain Imaging Lab and UW-Milwaukee Neuroscience Area for their feedback on this work.

