## Supplementary Materials for "Adult lifespan effects on functional specialization along the hippocampal long axis"

Supplemental materials

| Table S1 | | |
| --- | --- | --- |
| *List of cortical and subcortical abbreviations, full name, and Freesurfer label* | | |
| Abbreviation in Figure 4 | Full name | Freesurfer label |
| Frontal pole | Frontal pole | ctx-lh-frontalpole |
| VMPFC | Ventral medial prefrontal cortex | ctx-rh-medialorbitofrontal |
| Lateral OFC | Lateral orbitofrontal cortex | ctx-rh-lateralorbitofrontal |
| IFG (BA 44) | Inferior frontal gyrus – pars opercularis | ctx-rh-parsopercularis |
| IFG (BA 47) | Inferior frontal gyrus – pars orbitalis | ctx-rh-parsorbitalis |
| IFG (BA 45) | Inferior frontal gyrus –  pars triangularis | ctx-rh-parstriangularis |
| Rostral MFG | Rostral middle frontal cortex | ctx-rh-rostralmiddlefrontal |
| Caudal MFG | Caudal middle frontal cortex | ctx-rh-caudalmiddlefrontal |
| SFG | Superior frontal gyrus | ctx-rh-superiorfrontal |
| ACC | Rostral anterior cingulate cortex | ctx-rh-rostralanteriorcingulate |
| Mid cingulate | Caudal midcingulate cortex | ctx-rh-caudalanteriorcingulate |
| Precentral | Precentral gyrus | ctx-rh-precentral |
| Insula | Insular cortex | ctx-rh-insula |
| Temp. pole | Temporal pole | ctx-rh-temporalpole |
| Trans. Temp. | Transverse temporal gyrus | ctx-rh-transversetemporal |
| STG | Superior temporal gyrus | ctx-rh-superiortemporal |
| STS | Banks of the superior temporal sulcus | ctx-lh-bankssts |
| MTG | Middle temporal gyrus | ctx-lh-middletemporal |
| ITG | Inferior temporal gyrus | ctx-lh-inferiortemporal |
| Entorhinal | Entorhinal cortex | ctx-lh-entorhinal |
| PHG | Parahippocampal gyrus | ctx-lh-parahippocampal |
| Postcentral | Postcentral gyrus | ctx-lh-postcentral |
| Paracentral | Paracentral lobule | ctx-lh-paracentral |
| Precuneus | Precuneus | ctx-lh-precuneus |
| Sup. Parietal | Superior parietal lobule | ctx-lh-superiorparietal |
| Supramarginal | Supramarginal gyrus | ctx-lh-supramarginal |
| Inf. Parietal | Inferior parietal lobule | ctx-lh-inferiorparietal |
| PCC | Posterior cingulate cortex | ctx-lh-posteriorcingulate |
| Retrosplenial | Isthmus of cingulate gyrus | ctx-lh-isthmuscingulate |
| Fusiform | Fusiform gyrus | ctx-lh-fusiform |
| LOC | Lateral occipital cortex | ctx-lh-lateraloccipital |
| Lingual | Lingual gyrus | ctx-lh-lingual |
| Pericalcarine | Pericalcarine cortex | ctx-lh-pericalcarine |
| Cuneus | Cuneus | ctx-lh-cuneus |
| Caudate | Caudate nucleus | Left-caudate |
| Putamen | Putamen | Left-Putamen |
| Pallidum | Globus pallidus | Left-Pallidum |
| Thalamus | Thalamus | Left-Thalamus |
| Amygdala | Amygdala | Left-Amygdala |
| NAC | Nucleus accumbens area | Left-Accumbens-area |
| Ventral DC | Ventral diencephalon | Left-VentralDC |
| Cerebellum | Cerebellum | Left-Cerebellum-Cortex |

| Table S2 | | | | | | | | | |
| --- | --- | --- | --- | --- | --- | --- | --- | --- | --- |
| *Coefficients (p-values) from models using connectivity to predict episodic memory* | | | | | | | | | |
|  | Tail | | | Body | | | Head | | |
| Target Region | F | Y | O | F | Y | O | F | Y | O |
| Frontal pole | -- | -- | -- | -- | -- | -- | -- | -1.19 (.12) | -- |
| VMPFC | .24 (.68) | -- | -- | -- | -- | -- | -- | -- | -- |
| Lateral OFC | -- | -- | -.12 (.90) | -- | -- | -1.50 (.11) | -- | -- | -- |
| IFG BA44 | .25 (.71) | -- | -- | -- | -- | -- | -- | -- | -- |
| IFG BA47 | -- | -- | -- | -- | 1.16 (.17) | -- | -- | .28 (.76) | -- |
| IFG BA45 | -- | -- | -.43 (.74) | -- | -- | -- | -- | -- | -- |
| Rostral MFG | -- | -- | .85 (.58) | -- | -.08 (.93) | -- | -- | -- | -- |
| Caudal MFG | -- | 1.76 (.06) | -- | -- | -- | -- | -- | -- | -- |
| ACC | -- | -- | -- | -.54 (.28) | -- | -- | .19 (.71) | -- | -- |
| Mid cingulate | -.17 (.75) | -- | -- | -- | -.72 (.35) | -.09 (.95) | -- | -1.52 (.10) | **1.66 (.01)** |
| Precentral | -- | -- | -- | -- | -.98 (.36) | -- | -- | -- | -- |
| Insula | -- | -- | .58 (.64) | -- | -- | -- | -- | -- | -- |
| Temporal pole | -- | -- | -- | .41 (.47) | -- | **2.05 (.01)** | -- | -- | -- |
| Transverse temporal | -- | -- | -.93 (.21) | -- | 1.34 (.18) | -- | .98 (.08) | 1.72 (.17) | -- |
| STG | -- | -- | -- | -- | -- | -- | -.76 (.18) | -- | -- |
| STS | -.44 (.52) | -1.07 (.37) | -- | -- | .61 (.62) | -- | -- | -- | -- |
| MTG | -- | -.51 (.65) | -- | -- | -.52 (.57) | -.39 (.73) | -- | **-1.97 (.048)** | .71 (.58) |
| Entorhinal | -- | .66 (.48) | -- | -- | -- | -- | -- | -- | -- |
| PHG | -- | -2.10 (.05) | -- | -- | **1.80 (.03)** | .40 (.64) | -- | **-2.52 (.001)** | .65 (.36) |
| Paracentral | -- | -- | -- | -- | -.68 (.37) | .36 (.67) | -- | -- | -1.54 (.13) |
| Precuneus | -- | -- | -- | -- | -- | -- | -- | 1.27 (.28) | -- |
| Superior parietal | **1.60 (.02)** | 1.17 (.16) | -- | -- | -- | -- | -- | 1.21 (.16) | -- |
| Inferior parietal | -- | -- | -- | .13 (.80) | -- | -- | -- | .58 (.42) | -- |
| Retrosplenial | -- | -- | -- | -- | -- | -- | .41 (.50) | -.01 (.99) | -- |
| Fusiform | -.83 (.07) | -- | -.83 (.50) | -- | -.97 (.13) | .54 (.55) | -- | -- | -- |
| Lingual | .25 (.65) | -- | -- | -- | -- | .52 (.61) | -- | -- | .87 (.25) |
| Pericalcarine | -- | -- | -- | -- | -- | -- | -- | -1.38 (.06) | -- |
| Cuneus | -- | -- | -- | -- | .54 (.50) | -- | -- | -- | -- |
| Caudate | -- | 1.13 (.20) | -- | -- | -- | -- | -- | -- | -- |
| Putamen | -.60 (.48) | -- | -1.17 (.30) | .07 (.93) | -1.38 (.24) | -- | -.46 (.47) | -- | -- |
| Pallidum | -- | .46 (.72) | -- | -- | .92 (.48) | -- | -- | -- | **-3.05 (.007)** |
| Thalamus | -- | -- | -- | -- | -- | -- | -- | -.26 (.70) | -- |
| Amygdala | -- | .75 (.53) | -- | -- | -- | -- | -- | -- | **1.46 (.046)** |
| NAC | -- | -- | -- | -- | -- | -- | -- | -- | **2.42 (.002)** |
| Ventral DC | **1.94 (<.001)** | .42 (.73) | **2.41 (.01)** | -- | .99 (.22) | -- | -- | -- | -- |
| Cerebellum | -- | .22 (.80) | -.85 (.36) | -- | -- | .54 (.71) | -- | .29 (.69) | -1.43 (.13) |

*Note:* F = model from full sample, Y = model based on younger half of the sample, O = model based on older half of the sample; -- represents connections that were not included in the final model; bolded connections were significant predictors at an uncorrected alpha = .05; target ROIs that are not listed had no connections that were included in the final model.
